# A conserved transcriptional backbone and rewiring of gene-regulatory networks in activated human CD4⁺ T cells

**DOI:** 10.1101/2025.11.25.687998

**Authors:** Ionut Sebastian Mihai, Iker Núñez-Carpintero, Davide Cirillo, Johan Trygg, Mattias Forsell, Martin Selinger

**Author notes:** These authors contributed equally to this work (Ionut Sebastian Mihai, Martin Selinger). **Contact:** Ionut Sebastian Mihai, Martin Selinger, Iker Núñez Carpintero, Davide Cirillo, Johan Trygg, Mattias Forsell.

## Abstract

CD4+ T cells are components of the adaptive immune system with a plethora of subtype-specific functions. In order to further dissect the activation and differentiation regulatory program(s) of individual CD4+ T cell subsets, we performed an in vitro activation and differentiation of human primary naive CD4+ T cells towards Th1, Th2, Th17 and Treg subtypes followed by the single-cell RNA-seq and ATAC-seq (multiome) analysis. Resulting multiome data were used for constructing the subtype-specific gene regulatory networks, which were next assessed for their differences/similarities among the subtypes. Surprisingly, a conserved set of 8 “backbone” transcription factors (TFs) was identified as highly central in all subtypes, however, with unique differentiation-driven rewiring tendency. Subtype-specific “driver” TFs were identified in the case of Th1-Th1_17-Th17 lineage (*EOMES*, *HLF*), naive Tregs (*ESR1*, *DACH1*), and memory Tregs (*SOX13*). Finally, we applied community detection algorithms to identify potential non-obvious groups of genes that regulate diverse molecular functions within the differentiated subtypes, linked to the backbone TFs. Our atlas aims at providing a high resolution understanding of the gene regulatory networks and their rewiring in human primary CD4+ T cells, upon activation and differentiation.

## Introduction

CD4+ T cells are key components of the adaptive immune system which carry out a wide variety of actions: from recruiting and activating other cellular components of the immune system to the site of infection, to fine-tuning the immune response. In order to transition from a naive to a functional differentiated state, a large number of environmental cues are required to decide on a suitable state - referred to as T cell ‘subtypes’. Commonly differentiated subtypes from naïve CD4+ T cells include (but are not restricted to) Th1, Th2, Th17 and Treg. These commonly studied subtypes are classified by either relative expression of canonical transcription factor (TF) markers (i.e TBX21, GATA3, RORC and FOXP3), by the secreted cytokine profile (i.e: IFNG, IL4, IL17 etc..) or by surface marker expression (i.e: CCR7, CD62L, IL2R etc.). Single-cell studies^1,2^ have further shed light on the heterogeneity among CD4+ T cells and that unbiased genomic characterization of the T cell states revealed the stochastic and complex picture to traditional CD4+ T cell subtypes. Furthermore, these studies point out a highly context-dependent rewiring pattern of TF activity, in terms of interaction partners and downstream regulation, in some cases referred to as “efectorness”^2^. One limitation however, is the lack of high resolution unbiased data and computational approaches to further investigate the rewiring. Gene regulatory networks (GRNs) are powerful tools for description and prediction of multiple biological events, depending on the network modeling process and the type of data used for network inference^3^. The emerging use of single-cell multiomics data, integrating RNA-seq and ATAC-seq data from the same cell, for GRN construction ^4,5^, poses as a higher resolution strategy for achieving a better understanding in the rewiring of genes under specific cellular polarizing conditions. Less common, yet suggested^6^, is the use of community-detection algorithms for exploration of the communities that compose the networks. Communities can be defined as a collection of nodes that are highly interconnected as a group within the GRN^7^. GRN community detection coupled with functional enrichment analysis on those communities, can help reveal the underlying molecular programs, their orchestrators as well as the stochastic and dynamic community composition behind CD4+ T cell activation and differentiation of different states. Despite this strategy not being widely common within the field, it has been previously reported in the context of cancer research^8,9^

Biological regulatory networks are inherently stochastic systems, where transcription factor activation, chromatin accessibility, and downstream gene expression fluctuate due to intrinsic biochemical noise and extrinsic environmental variability^10,11^. In such systems, information does not propagate deterministically along regulatory edges, but instead diffuses probabilistically through the network, with transitions between states better described by Markovian dynamics than by static deterministic rules. This stochastic diffusion perspective aligns naturally with random-walk–based community detection, where communities emerge as regions of high retention probability for simulated walkers moving across the graph^12,13^. In gene regulatory networks, such random-walk retention corresponds to regulatory “basins” or locally stable modules, capturing how noisy TF signals preferentially circulate within functional subcircuits rather than across the entire network. Foundational work in network science has demonstrated that random-walk and Markov-stability approaches outperform purely topological clustering in recovering biologically meaningful modules in complex networks^12,13,14^, supporting their application for identifying regulatory programs in heterogeneous single-cell datasets.

Recent multiomic studies have begun to characterize chromatin-linked regulatory programs in human lymphocytes and connect them to disease-associated variants, but a network-level understanding of how human primary CD4⁺ T cells reorganize their regulatory circuitry across activation and lineage polarization remains limited^15,16,17^. Existing atlases primarily focus on transcriptional trajectories or enhancer activity, with less emphasis on quantifying (i) which TFs form a conserved “activation backbone”, (ii) which TF connectivity patterns are selectively rewired in specific helper lineages, and (iii) how regulatory communities reorganize across these states.

To better understand the various CD4+ T cell states, and which transcriptional networks orchestrate their differentiation program, we have used a combined single-cell RNA-seq and ATAC-seq protocol. Independent use of single RNA-seq and ATAC-seq has successfully delineated the trajectories of, e.g., the development of αβ T cells^18^. By applying a paired multiomic readout that directly links chromatin accessibility to transcriptomic output in each cell, and leveraging the integrative power of GRN construction and graph-theoretic community detection, we provide a systems-level view of regulatory rewiring in activated and differentiated human CD4⁺ T cells. Through this approach, we identify conserved regulatory “backbone” TFs, lineage-biased “driver” TFs, and condition-specific changes in community architecture. All-in-all, our atlas points toward a systemic view of CD4+ T cell states, relevant for adoptive cell therapy and bioreactor design.

## Results

### Multiome analysis of human *in vitro* activated and differentiated CD4+ T cells unveiled canonical Th subtypes and CD4-/CD3+ NKT cells

In order to study the regulatory networks responsible for the activation of naïve CD4+ T cells and their differentiation into specific subtypes, a combined scRNA-seq and scATAC-seq (multiome) analysis of human Th1-, Th2-, Th17-, and Treg-differentiated cells was performed. For this purpose, we established an *in vitro* differentiation model (Fig. 1A) where PBMCs from blood of healthy male donors were used for the isolation of naïve CD4+ T cells, which were anti-CD3/28 activated and simultaneously differentiated under Th1-, Th2-, Th17-, and Treg-polarizing conditions using particular cytokine/antibody mix for 5 days (Table S1). To assess the differentiation efficiency, we first analyzed the expression of Th1/Th2/Th17/Treg canonical markers TBX21, GATA3, RORC, and FOXP3, respectively. In all cases, the population of differentiated cells showed increased levels of the respective canonical marker when compared to naïve non-activated cells (Fig. 1B, Fig. S1A).

**Figure 1:**
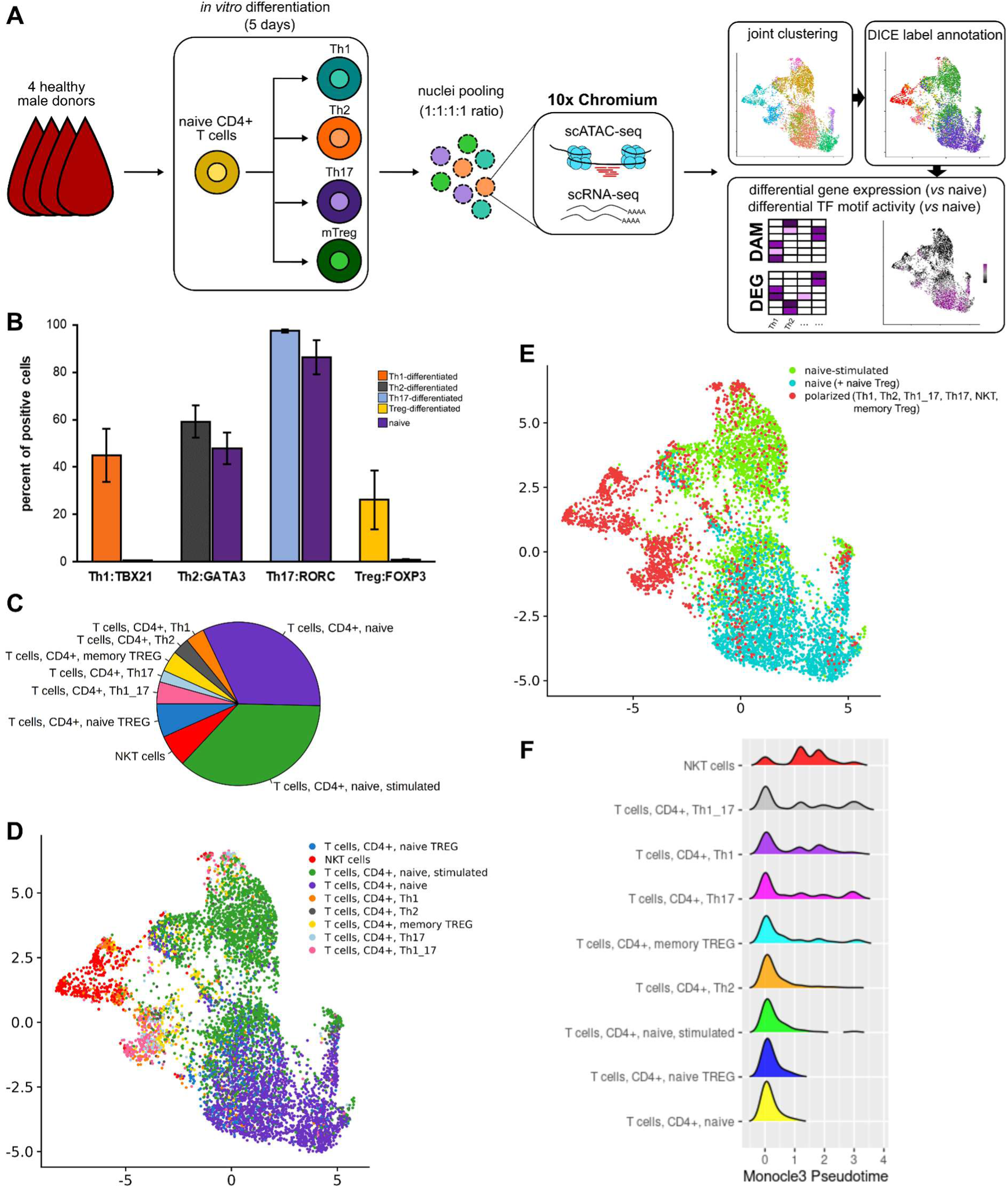
Multiome analysis of human in vitro activated and differentiated CD4+ T cells. (A) Experimental workflow for multiome-seq analysis. (B) Human naive CD4+ T cells were activated and differentiated using Th1-/Th2-/Th17/Treg-specific cytokine/antibody cocktails for 5 days. Differentiated populations were analyzed for the expression of canonical markers (TBX21, GATA3, RORC, FOXP3) using FACS detection of antibody-labeled cells. Values are expressed as the mean percentage (±SD) of marker-positive cells isolated from three donors. (C) The distribution of DICE-annotated T cell subtypes within the dataset. (D) Joined (scRNA-seq + scATAC-seq) UMAP embedding with cells highlighted according to the DICE annotation of T cell subtypes (left) or (E) activation/differentiation state. (F) Pseudotime analysis of cell trajectories.

Prior to the multiome analysis, cells were first sorted for high viability (all sizes of viable cells were included to capture the differentiation dynamics). Afterwards, the nuclei of Th1-, Th2-, Th17-, and Treg-polarized CD4+ T cells were extracted, pooled in equal ratios, and processed on the 10x Genomics Chromium platform (Fig.1A). After initial pre-processing of scATAC-seq and scRNA-seq data using Seurat and Signac packages^19,20^, 7069 cells were used for further analysis. We first performed Uniform Manifold Approximation and Projection (UMAP)^21^ for scRNA-seq followed by the cell type annotation using SingleR^22^ and DICE database of immune cells^23^ (Fig. S2A). Automatic annotation resulted in identification of 9 T cell subtypes, where the vast majority was annotated as CD4+ T naïve (32.4%) and CD4+ T naïve-stimulated (36.6%) cells. Abundancy of terminally differentiated CD4+ T cell subtypes of Th1, Th2, Th1_17, Th17, and Treg memory cells was ranging between 2.3 – 4.3% (Fig. 1C). To validate the results of DICE-linked annotation, the expression pattern of canonical marker genes for each subtype was verified manually (Fig. S2B). A subset of 453 cells (6.4%) was annotated by SingleR as NK cells (CD56/NCAM1+, CD161/KLRB1+, CD94/KLRD1+, GNLY+, NKG7+). However, based on their CD3+ phenotype (Fig. S2C), we re-annotated this subset as NKT cells^24^. To determine the origin of NKT cells in our dataset and verify their CD3/CD4 status, we analyzed for their presence in isolated naive CD4+ T cells by the detection of an invariant Vα24-Jα18 TCRα receptor^25^ together with CD3 and CD4. Interestingly, 5% of CD4+ isolated cells were documented as Vα24-Jα18 positive, with the vast majority of them (91%) being also double-positive for both CD3 and CD4 (Fig S1B).

The cell type labels annotated in the scRNA-seq dataset were subsequently transferred to the scATAC-seq UMAP, where those followed a very similar clustering pattern (Fig. S2B).The joint UMAP visualization combining the features from both gene expression and DNA accessibility, illustrates co-clustering of (i) terminally differentiated subtypes, mainly NKT cells and Th1/Th17/Th1_17 cells, and (ii) naïve/naïve-stimulated T cells/naïve Tregs (Fig. 1D). In order to further describe the relationships between individual T cell subsets, we constructed phylogenetic trees using PCA (scRNA-seq) and LSI (scATAC-seq) dimensional reductions (Fig. S2D). Interestingly, NKT cells clustered as an outgroup in both cases, suggesting their distinctive transcriptomic and epigenetic profile. Similar clustering outcome was observed also in the case of Th1_17/Th17 subsets. Overall, the comparison of both cladograms documents a very high correlation between the epigenetic status and canonical functional annotation of CD4+ T cell subsets.

### Comparison between naive and activated CD4+ T cell subsets reveals conserved and subtype-specific features

To verify the DICE-derived cell annotation and also identify novel T cell subtype-specific genes (scRNA-seq) and open TF motifs (scATAC-seq), we searched for differentially expressed (DE) genes and differentially active (DA) TF motifs using Seurat^19^ and ChromVAR^26^ packages, respectively. For both analyses, CD4+ T naïve cells were used as a reference, so the activation/differentiation- and naïve-linked factors could be identified and compared between subtypes to identify not only the subtype-specific marker genes/motifs, but also general markers for activation and naiveness.

In total, 2,730 DE genes were identified (p<0.01, log_2_-FC > 0.25) among all comparisons; the Szymkiewicz-Simpson overlap index (SSI) values ranged between 0.25 to 0.90, with the highest overlap ratios calculated for Th2 and Treg-memory cells, followed by Th2 and naïve-stimulated cells (Fig. 2A). Importantly, hierarchical clustering of T cell subsets based on SSI values closely followed the phylogenetic clustering for scRNA-seq data (Fig. 1E) - the differentiation axis of Th1-Th1_17-Th17 subsets co-clustered together with NKT cells, whereas Th2 clustered within the group of Tregs and naïve-activated cells. The SSI values for up-regulated DA motifs (p<0.01, log_2_-FC > 0.25) were substantially higher with values ranging between 0.82 and 1.00 (Fig. 2A), indicating the presence of highly conserved TFs involved in CD4+ T cell activation. Similarly to DE genes, the highest SSI value for DA motifs was identified for Th2 and naïve-stimulated cells. Highly correlative results were described also in case of clustering, where Th1 and Th1_17 subsets co-clustered with NKT cells whereas Th2 cells co-clustered with Tregs and naïve-activated cells (Fig. 2A).

**Figure 2:**
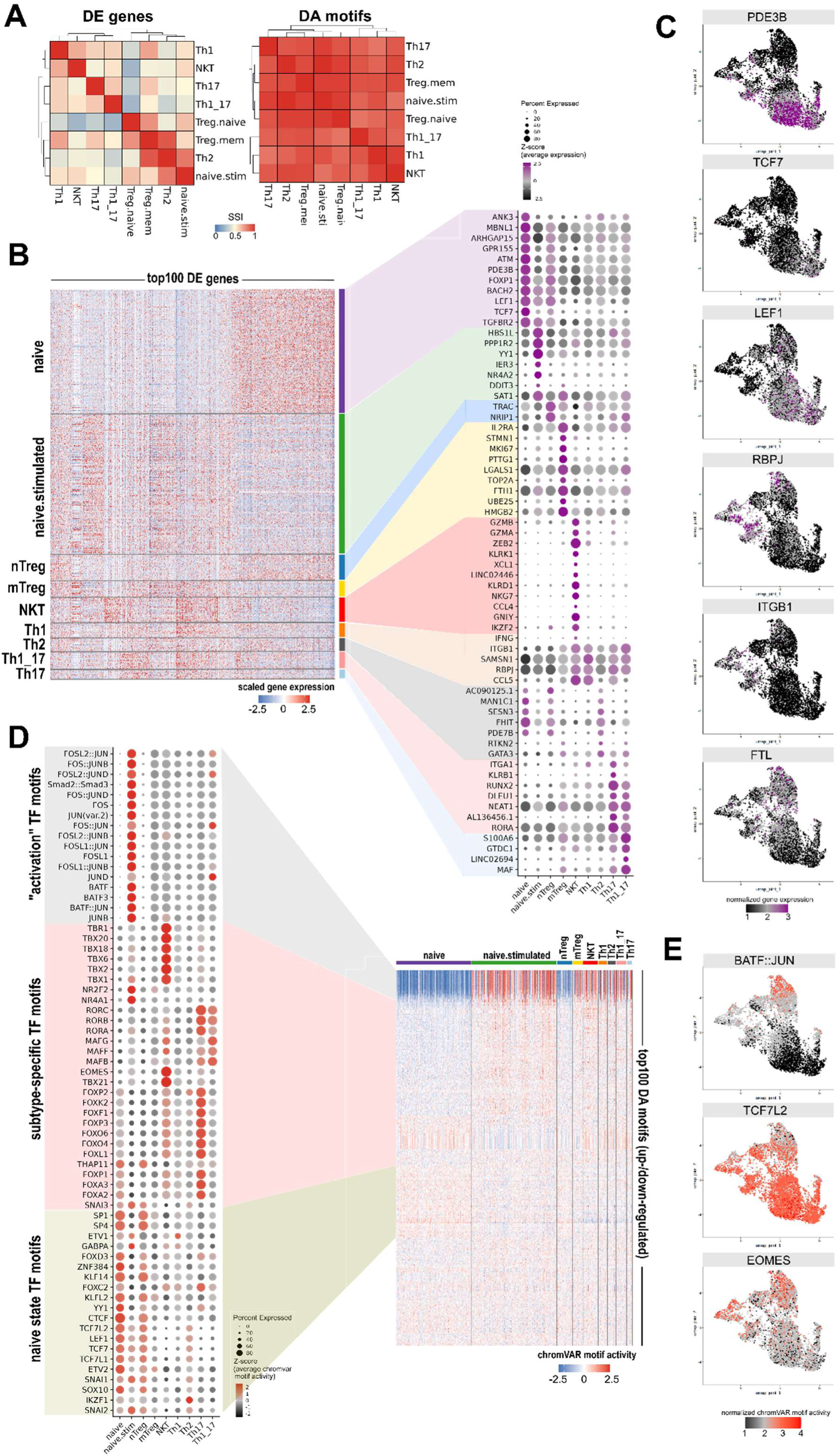
Identification of marker genes and TF motifs in DICE-annotated CD4+ T cell subsets. (A) Heatmap representing Szymkiewicz-Simpson overlap coefficient (SSI) values for DE genes (left) and DA TF motifs (right) among T cell subsets, identified by Seurat and chromvar/Signac, respectively. (B) Heatmap representing scaled single-cell expression of top 100-ranked (up/down) DE genes from all T cell subsets; dot plot illustrates the Z-score of average expression of selected marker genes. (C) UMAP-embedded visualization of selected marker gene expression. (D) Heatmap representing normalized single-cell TF motif activity (chromVAR) of top 100-ranked (up/down) DA motifs from all T cell subsets; dot plot illustrates the Z-score of average activity of selected marker motifs. (E) UMAP-embedded visualization of selected marker TF motifs calculated by chromVAR.

The visualization of single-cell expression through a heatmap for the top 100 ranked DE genes reveals distinct clusters of marker genes specific to different T cell subtypes (Fig. 2B). These findings highlight the heterogeneity and unique gene expression patterns associated with each T cell subtype, as previously described by Cano Gamez et al.^2^. Furthermore, we conducted a comprehensive search for general activation markers, i.e. genes that were identified as DE across all activated T cell subsets. We identified 50 genes to be up-regulated across all the activated cell types with RBPJ, ITGB1, MAF, LGALS1, GTDC1, TNFRSF4, and FTL as the top-ranked genes (Table S2; Fig. 2C). In order to further explore the functional differences among activated T cell subsets, the gene set enrichment analysis (GSEA) of identified DE genes was performed. Significantly enriched GO terms were identified only in the case of NKTs, CD4+ T naive-stimulated, and memory Treg subsets (Fig. S2E). These were found to further support the DICE-based annotation, especially in the case of NKT cells with response to IFN-γ and NK cell mediated cytotoxicity among the top-scored terms. In case of scATAC-seq data, all activated subsets shared the very same top-scoring DA motif features recognized by the AP-1 family of TFs, including ATF subfamily (BATF, BATF3), JUN subfamily (JUN, JUNB, JUND), and FOS subfamily (FOS, FOSL1, FOSL2) (Fig. 2D). The AP-1 family of dimeric TFs regulates a plethora of cell processes and was shown to have an important role in activation and differentiation of CD4+ T cells as well^27^. Except for the AP-1 family, motifs recognized by the SMAD family of TFs (SMAD2, SMAD3) were also identified as top-ranked DA motifs. SMAD family TFs were described to be involved in TGF-β signaling, a key player in maintaining the T cell homeostasis and CD4+ T cell activation^28^. Apart from top-ranked “activation motifs”, we further identified “naive state motifs” and motifs specific for one or more T cell subtypes (Fig. 2D-E).

In case of Th1 cells, no high-ranked DE genes have been found to be expressed predominantly within this subtype only. Instead, the vast majority of Th1 DE genes was documented to be highly up-regulated also in Th1_17, Th17, or NKT cells, e.g. CCL5, RBPJ, ZEB2, SAMSN1, and ITGB1 (Fig 2B). Apart from those, an elevated expression of IFN-γ (IFNG), a signature cytokine for Th1 and NK/NKT cells^29^, was detected in part of Th1 (14% cells) and NKT cells (20% cells) (Fig. 2B). Similar situation of rather shared than Th1-unique features was also observed in case of DA motifs. The top-ranking Th1 DA motifs comprised of AP-1 and SMAD family TFs, which were highly active in all cell types (Fig. 2D). Additionally, the motif of the Th1 canonical marker^30^ TBX21, showed high activity scores in the Th1 subset, but also in Th1_17 and NKT cells (Fig. 2D). Interestingly, the EOMES-binding motif also showed high activity in Th1, Th1_17, and NKT cells. EOMES is linked to the cytotoxic effector phenotype in CD8+ and NK cells^31^ and recently was also shown to orchestrate the transition from Th17 to non-classical Th1^32^, thus highlighting the transition state of Th1_17 subset. Moreover, EOMES was also recently shown to be a key factor required for iNKTdevelopment^33^, which supports its high activity in the NKT subtype.

Similarly to Th1, no specific markers for Th2 cells have been identified, especially due to the high overlap values with memory Tregs and naïve-stimulated T cells (SSI values 0.90 and 0.82, respectively; Fig. 2A). We identified only two DE genes, GATA3 and RTKN2, having the highest expression in the Th2 subset (Fig. 2B). However, both genes were highly expressed also in Th17 and memory Tregs. Additionally, Th2 cells share a set of genes highly expressed in naïve and naïve Treg cells (e.g. PDE7B, FHIT, SESN3, CYSTLTR1, MAN1C1, AC090125.1). None of the DA motifs was found to be Th2-specific (Fig. 2D).

Marker genes shared between Th1_17 and Th17 subsets include RORA and 3 lncRNAs (AL136456.1, NEAT1, and DLEU1), out of which NEAT1 was shown to be crucial for Th17 differentiation process^34^. Elevated expression in the case of Th1_17, but not Th17, was documented for CCL5, GZMB, RUNX2, KLRB1, and ITGA1 genes, which were also highly expressed in Th1 and NKT cells. Vice versa, DE genes with higher expression in Th17 cells encompass MAF, LINC02694, GTDC1, and S100A6 (Fig. 2B). Except for canonical Th17 TF RORC^35^, ChromVAR analysis further identified MAFB, MAFF, MAFG, RORA, and RORB as Th1_17/Th17-specific DA motifs (Fig. 2D).

Memory Tregs showed a distinctive expression pattern in the case of HMGB2, UBE2S, FTH1, TOP2A, LGALS1, PTTG1, MKI67, and STMN1 genes. Out of these HMGB2 and MKI67 were recently linked to the effector Treg differentiation process^36,37^. Alpha subunit of IL-2 receptor (IL2RA) was documented to have the highest expression in memory Tregs as well, however, considerably high levels of IL2RA were documented in other subtypes such as Th1_17 and Th17 cells (Fig. 2B). None of the DA motifs was found to be specifically more active in memory Tregs.

Cells annotated as CD4+ naïve-stimulated represent an early state of activation^23^, which can be also considered as an intermediary transition state between naive and effector phenotype. The transient state of naïve-stimulated cells is documented by the list of DE genes and DA motifs, where most of these were identified as up-regulated in effector subsets, and simultaneously, by the retained expression of several genes highly expressed in naïve T cells. The top-ranking DE activation genes for naïve-stimulated T cells are SAT1 and DDIT3, which have similar fold-change also in memory Tregs and Th17 cells. Specific DE activation genes for naïve-stimulated cells then include NR4A2, IER3, YY1, PPP1R2 or HBS1L (Fig. 2B). On the opposite side, genes linked to the naïve state, e.g. LEF1, TGFBR2 or BACH2^38^, were still highly expressed in naïve-stimulated cells as well. Analogous situation was described for the motifs, where naïve-linked motifs for LEF1, TCF7, and TCF7L1-2 were found to be still highly active (Fig. 2D). In order to emphasize this ambiguity in terms of shared naive/effector motif activity, we visualized the normalized ATAC counts for top-ranked naive DA motif TCF7L2 and top-ranked activation motif BATF::JUN (Fig. 2E). Additionally, motifs for NR4A1 and NR2F2 were found to be the most up-regulated DA features specific for naïve-stimulated subset (Fig. 2D). These findings correspond with the early activation state of naïve-stimulated T cells, since the NR4A family of TFs and IER3 were shown to be involved in downstream signaling of activated TCR^39,40^.

Joint UMAP illustrates that naïve Tregs cluster on the transition border between naïve T cells and naïve-stimulated T cells, suggesting a close relationship. Indeed, the second highest high SSI value (0.64) for naïve Tregs was described in overlap with naïve-stimulated cells. The highest SSI value (0.75) was determined between naïve and memory Tregs (Fig. 2A). Based on the comparisons of DE genes with other cell subsets, NRIP1 and TRAC genes were identified as naïve Treg-specific. Moreover, high expression of IKZF2 was described, a TF required for the inhibitory function of Tregs^41^. However, IKZF2 showed the highest expression value in NKT cells (Fig. 2B). Naive Tregs in comparison to memory Tregs express low levels of IL2RA^42^, which was documented also in our dataset (Fig. 2B). No specific DA motifs have been found in case of naïve Tregs, nevertheless, the overall activity of “activation” motifs was significantly lower than in other subsets. Notably, naïve Tregs were also described to have an equally active naïve-state-linked motifs (e.g. CTCF, TCF7, LEF1, TCF7L1-2) when compared to the naïve T cells (Fig. 2D).

NKT cells represent a distinct population of CD1d-restricted T cells that share several phenotypic and functional characteristics with NK cells^24^. The subset of CD4-/CD3+ NKT cells forms a well-defined cluster in joint UMAP (Fig. 1D) with the most distinctive differentiation pattern from all annotated subsets. Marker DE genes specific to NKT cells only include GNLY, CCL4, NKG7, KLRD1, LINC02446 lncRNA, XCL1, and KLRK1. Several top-ranked DE genes in NKT cells (e.g. ZEB2, GZMA or GZMB) were shown to be strongly up-regulated also in the Th1-Th1_17-Th17 axis. In case of DA motifs, NKT cells share the high-ranked features with other subtypes, nevertheless, specific NKT motifs were identified as well. These comprise of EOMES, TBX family (TBX1/2/6/18/20/21) and TBR1. Importantly, all NKT-specific DA motifs were significantly upregulated in Th1, Th1_17, and naïve-stimulated T cells as well, albeit to a lesser extent.

Comparison of downregulated DE genes and DA motifs among subsets revealed a surprisingly conserved group of genes/motifs specific for the naïve state of CD4+ T cells. These include well-known TFs already linked to the naïve state such as TCF7^43,44^, LEF1^43,44^, BACH2^38^ or FOXP1^45^, however, novel genes were identified as well including PDE3B, ATM, GPR155, ARHGAP15, MBNL1, and ANK3 (Fig. 2B). The list of top-scoring DA motifs identified as characteristic for the naïve state further supports the annotation, when LEF1 and TCF7 binding motifs were identified together with CTCF, TCF7L1, TCF7L2, ETV2, and SOX10 motifs (Fig. 2D). As already mentioned above, a substantial portion of described naïve CD4 +T cell markers was associated with a characteristic pattern of high expression also in naïve Treg cells and naïve-stimulated cells.

### Analysis of CD4+ T cell GRNs reveals extensive rewiring processes upon activation and differentiation

Previous attempts have been made to elucidate the complex differentiation program of CD4+ T cells^46^, but lacked the high resolution of multiome technologies. Our CD4+ T cell multiome data with annotated T cell subsets provide such a resolution and thus, we constructed the GRN for each differentiated subtype individually, by making use of Pando^4^. As biological networks tend to be highly modular, we further investigated the biological relevance of highly-central nodes identified in each subtype’s GRN by performing the community detection (Fig. 3A).

**Figure 3:**
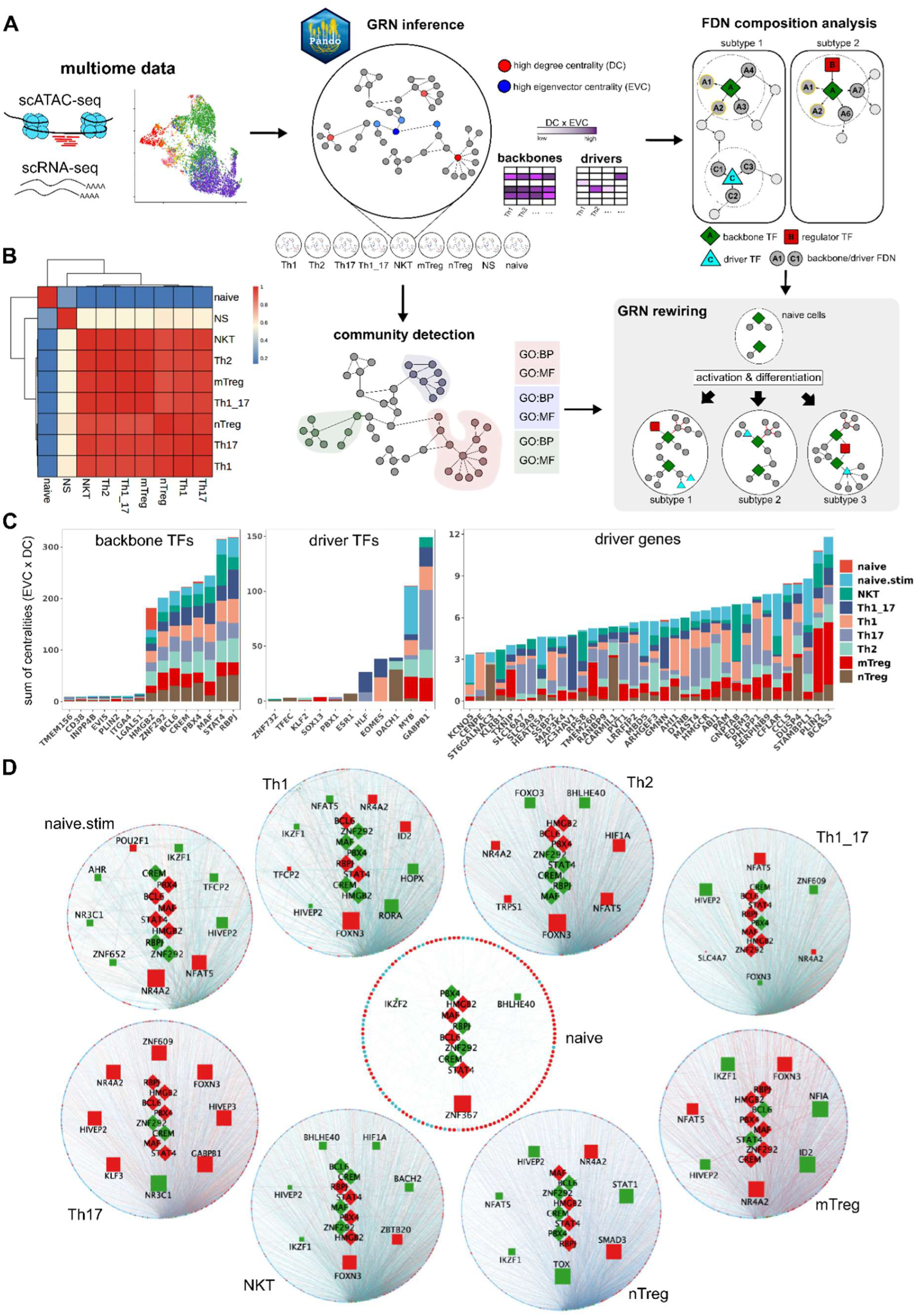
GRN analysis and TF backbone properties. (A) Experimental workflow for GRN analysis. (B) Jaccard indices describing the composition overlap of individual GRNs inferred by Pando. (C) Cumulative sum of “weighted” centrality as a result of combining the eigenvector centrality (EVC) and degree centrality (DC) for backbone TFs, driver TFs, and driver genes. (D) Overall representation of the networks, with top-highest centrality. Squares represent Backbone TF neighbors, whilst rhombi represent backbone genes. Red color represents target of backbone genes, whilst green represents regulators. In case of connections (edges), the red represents negative regulatory effect and green represents positive regulatory effect.

Overall, the GRNs derived from the CD4+ T cell effector (T_eff_) states had a high overlap in terms of node composition among themselves (minimum Jaccard = 0.93). On the contrary, the lowest overlapping values were shown for naive *vs* all remaining T_eff_ states (maximum Jaccard = 0.14), thus hinting at an increase in size and node composition, as a result of activation and differentiation cues. Interestingly, naive-stimulated cells exhibited intermediate overlap values for both Teff subtypes (maximum Jaccard = 0.56) and naive cells (Jaccard = 0.26), hence highlighting their transition state between naive and differentiated phenotype (Fig. 3B).

As a consequence of high modularity, biological networks tend to have a power-law degree distribution (i.e. the ranked total sum of connections is often “monopolized” by a discrete number of TFs)^47^. In order to identify such TFs we searched for genes with high degree centrality and high eigenvector centrality as proxy metrics. As a result of performing a node intersection of all the GRNs, two functional groups of highly central TFs were defined: (i) the backbone TFs having high degree centrality and high eigenvector centrality in all subtypes, irrespective of the activation and stimulation status, and (ii) the driver TFs having high degree centrality and high eigenvector centrality only in specific subtype(s). Furthermore, with regards to the backbone/driver TF position within the network, two additional gene categories were distinguished: (i) the regulators as first degree neighbors (FDNs) having targeting connections towards the backbone/driver TFs (including other backbone TFs), and (ii) targets as FDNs (including other backbone TFs) with incoming connections from the backbone/driver TFs (Fig. 3A).

The top-ranking backbone TFs include *BCL6, CREM, HMGB2, MAF, PBX4, RBPJ, STAT4* and *ZNF292*. Surprisingly, the eigenvector centrality and degree centrality for other TFs was significantly lower, which highlights the central role of backbone TFs within all the subtypes (Fig. 3C). Except for *PBX4* and *ZNF292*, all identified backbone TFs have already been associated with CD4+ T cell development^37,48–54^. These findings in combination with the visualization of backbone-centered GRNs for individual subtypes clearly demonstrates the conserved significance of backbone TFs (Fig. 3D). In the case of driver TFs, 11 genes were identified with *MYB* (naive-stimulated), *DACH1* (nTreg), *EOMES* (Th1, Th1_17), and *HLF* (Th17, Th1_17) as top-ranked ones (Table 1, Fig. 3C). Interestingly, 36 non-TF genes fulfilled the driver TF criteria as well, thus we created an additional group of driver genes (Table 1, Fig. 3C).

**Table 1:** Overview of identified driver TFs and driver genes.

| subtype | driver TFs | driver genes |
| --- | --- | --- |
| Th1 | EOMES | CCL5, CENPE, SERPINB9, PAM, SSBP2, AHI1, DTNB |
| Th2 | KLF2 | ABI1, ARHGEF3 |
| Th17 | HLF, GABPB1, PBX1 | PVT1 (lncRNA), HMGCR, MAST4, LRRFIP2, EDEM3, RANBP9, MAP3K4, SLC16A7, SLC9A9, PHLPP1 |
| Th1_17 | EOMES, HLF | GMNN, ZC3HAV1 |
| NKT | ZNF732 | RPS8, KLRB1, GNPTAB |
| mTreg | SOX13 | PLIN2, TXNIP, BCAS3, TMEM260 |
| nTreg | TFEC, DACH1, ESR1 | DUSP4, ST6GALNAC3, CARMIL1 |
| naive-stimulated | MYB | CFLAR, MBD5, STAMBPL1, HEATR5A, KCNQ5 |

The backbone-centered GRN visualization further suggests subtype-specific differences in terms of edge composition, distribution, and overall regulation hierarchy (Fig. 3D). Based on this observation combined with a subtype-distinctive pattern of DE genes and DA motifs, we hypothesized that, despite the high centrality of backbone TFs across all subtype GRNs, these are subject to an activation- and differentiation-induced network rewiring. The network rewiring is defined as both net change in number of interaction partners and change in their composition, which ultimately results in a functional switch within the backbone branch. To verify this, we investigated the size, composition, and functional annotation of backbone TFs’ FDNs across all the networks. To increase the functional resolution for individual backbone TFs, the analyses were performed separately on target FDNs and regulator FDNs. When comparing the number of target FDNs among individual networks, the subset of naive cells displayed the lowest amount of FDNs whereas all T_eff_ subsets exhibited considerably higher FDN numbers, which were comparably equal in size (Fig. 4A). When querying the composition overlaps of target FDNs and performing hierarchical clustering, obtained SSI values ranged between 0 to 0.69 with median value of 0.026 pointing towards low composition overlap in general (Fig. 4B). Nevertheless, distinct co-clustering of individual backbone TFs’ FDN subsets was observed for HMGB2 and RPBJ showing the highest SSI values (max SSI = 0.69 and 0.63, respectively) and forming complete clusters of all 9 cell subsets (Fig. 4C). Furthermore, a detailed HMGB2 target FDN composition analysis revealed 16 genes as “conserved” FDNs (i.e. genes present in more than 6 subtypes), whose biological function was found to be linked to the cell cycle regulation (p = 4.62E-10). Similar clustering and SSI values ranging between 0.10 to 0.45 were documented in the case of BCL6, CREM, and STAT4, however without the presence of naive FDN subsets within the formed clusters. On the contrary, extremely low overlap values among all cell subtypes were observed for MAF, PBX4, and ZNF292 (Fig. 4C). In fact, the subtype specificity of PBX4 target neighborhood is the most persistent across all cell subtypes of all backbone TFs, with a mean of 23.11 ± 4.42 subtype-specific target connections (SSTCs), which represents 54.03% of total SSTC number. Comparably, MAF and ZNF292 present similar rewiring dynamics showing a mean of 24.22 ± 8.76 (57.45%) and 22.00 ± 8.05 (48.21%) of SSCTs, respectively(Fig. S3A). These findings combined suggest that the rewiring is happening in a subtype-specific manner, where individual backbone TFs do not mostly share the downstream regulatory pathways.

**Figure 4:**
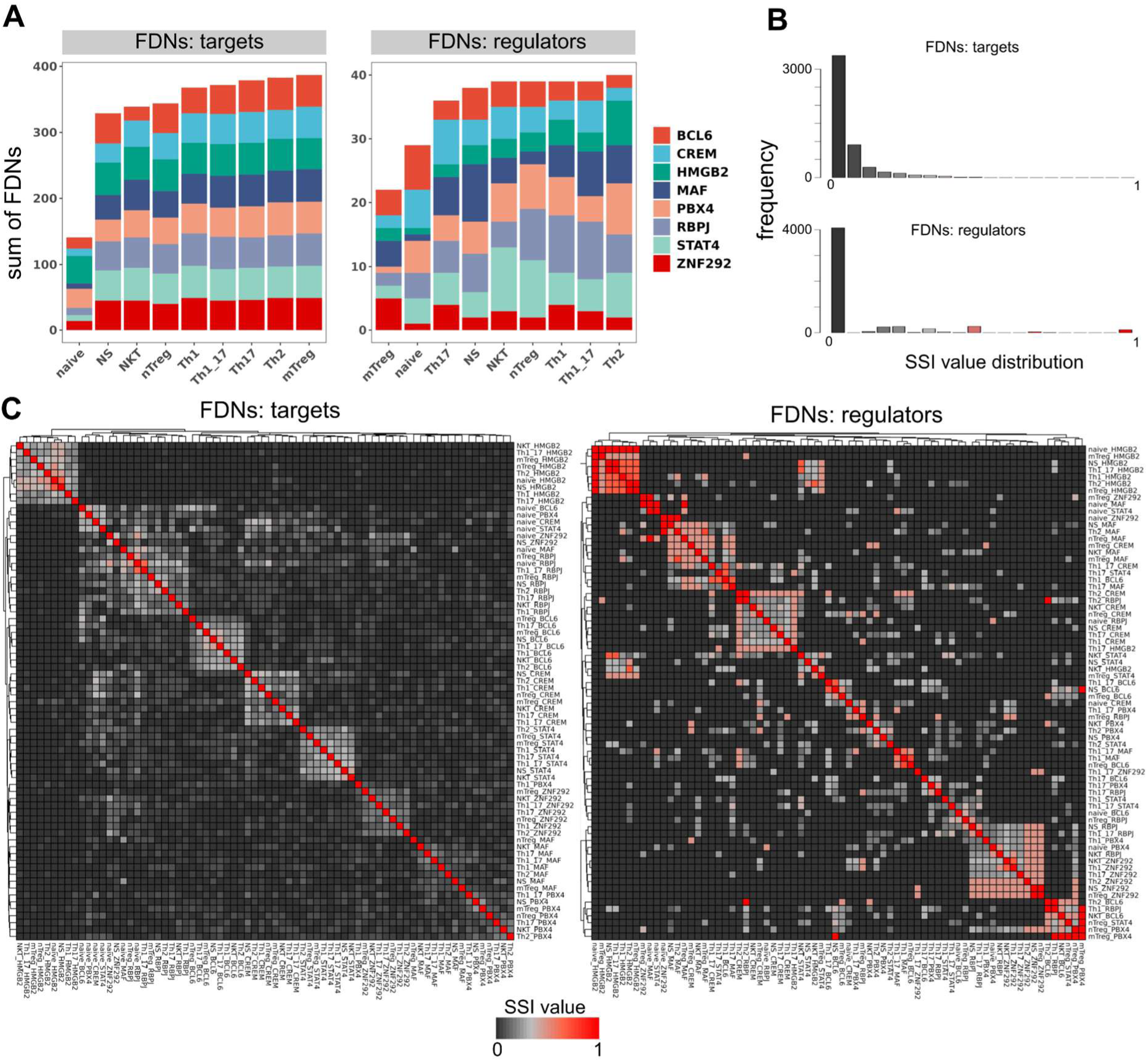
The first-degree neighborhood analysis of backbone TFs. (A) Cumulative sum of first-degree neighbors (FDNs) for backbone TFs - target genes (left) and regulator TFs (right). (B) Value frequency distribution of all Simpson-Szymkiewicz índices depicted in (C) Simpson-Szymkiewicz índices describing the composition overlap of FDN groups for individual backbone TFs - target genes (left) and regulator TFs (right).

While the subtype-specific differences in backbone TFs’ target FDN composition give a clear picture of ongoing rewiring processes, it is the subset of regulator TFs which might be responsible for activating these. Thus, we next focused on a detailed mapping of regulator TFs in each of the annotated T cell subtypes, and their edges. Overall, the numbers for identified regulator TFs were higher in all subsets than in naive cells, with the exception of mTregs (Fig. 4A). The composition of backbone TFs’ regulator FDNs revealed a conserved cluster of HMGB2 regulators (SSI values 0.5-1.0) in the case of all cell types except for NKT cells (Fig. 4C). Within this cluster *HMGB1* and *HIVEP2* were found as the most-conserved regulator TFs being present in 7 and 6 cell types, respectively. Similar clustering was documented for ZNF292 and CREM, however, with lower SSI values than in the case of HMGB2 (Fig. 4C). Notably, some of the backbone TFs exhibited the ability to act as either regulator or target of other backbone TFs (Fig. 3D).This phenomenon is especially important in the case of CREM and ZNF292, which were found to be highly-conserved regulators of each other in identical groups of 6 cell subtypes. Additionally, a conserved regulator TF *NFIA* (present in 6 cell subtypes) was identified for MAF, however, the rest of backbone TFs (PBX4, RBPJ, STAT4, BCL6) were absent of any conserved regulator TFs.

IKZF TF family members appeared amongst the top highly central regulator TFs in case of naive (IZKF2), and naive-stimulated, Th1, NKT, nTreg and mTreg (IKZF1) subtypes. Furthermore, HIVEP2 showed an overall regulator activity in the naive stimulated and all T_eff_ subtype networks, except for Th17, where it was a target. Further rewiring tendencies were observed for NFAT5, acting as regulator in Th1, and nTregs, whilst acting as a target in Th2, Th1_17 and mTreg. Similarly, FOXN3 acted as regulator solely for Th1_17 and Th17, whilst acting as target for Th1, Th2, NKT and memory Treg. Finally, some of FDNs were found to act exclusively as targets or regulators. This is the case of NR4A2 (target of all effector subtypes + naive stimulated) and BHLHE40 (regulator for naive, Th2 and NKT). Low SSI values in combination with very few conserved regulator TFs being identified further supports the dynamic rewiring of the backbone genes, not only at the downstream target level, but also upstream regulator level.

### Backbone-centric community analysis using random walk trap reveal network-specific molecular functions

Conceptually, biological networks are described as highly modular, containing sub-groups of densely-connected nodes, which are associated with specific biological functions. The number of potential biological processes occurring in a T cell that span beyond the first degree neighbors, at a given time point is still unknown. To uncover biologically relevant communities in the case of a particular CD4+ T cell subtype GSEA was performed on the communities identified by implementation of the random walktrap algorithm^55^. The enrichment was performed consulting the GO:MF and GO:BP databases and the resulting annotations are described in Table S3. We postulate that communities with at least 10 members, containing at least one backbone can yield useful insights on active molecular functions and further confirm the rewiring process. Thus, backbone TF-centric communities from all subtypes were analyzed for their composition overlap using SSI overlap values (Fig. 5A). We also searched for the presence of driver TFs and driver genes within these communities. The top-ranked driver TFs *EOMES* and *HLF* were not present in backbone TF-centric communities as well as *SOX13*, *ZNF732*, *KLF2*, and *TFEC*. In fact, *HLF, TFEC, and KLF2* were not found within any community (Fig. 5A, Table 2). Notably, we identified nTreg-derived community not only containing 5 out of 8 backbone TFs (*STAT4, RBPJ, PBX4, BCL6 and CREM*), but also *ESR1* and *DACH1* driver TFs. The robustness of this community suggests its high importance and relevance of *ESR1* and *DACH1* as potential key regulators.

**Figure 5:**
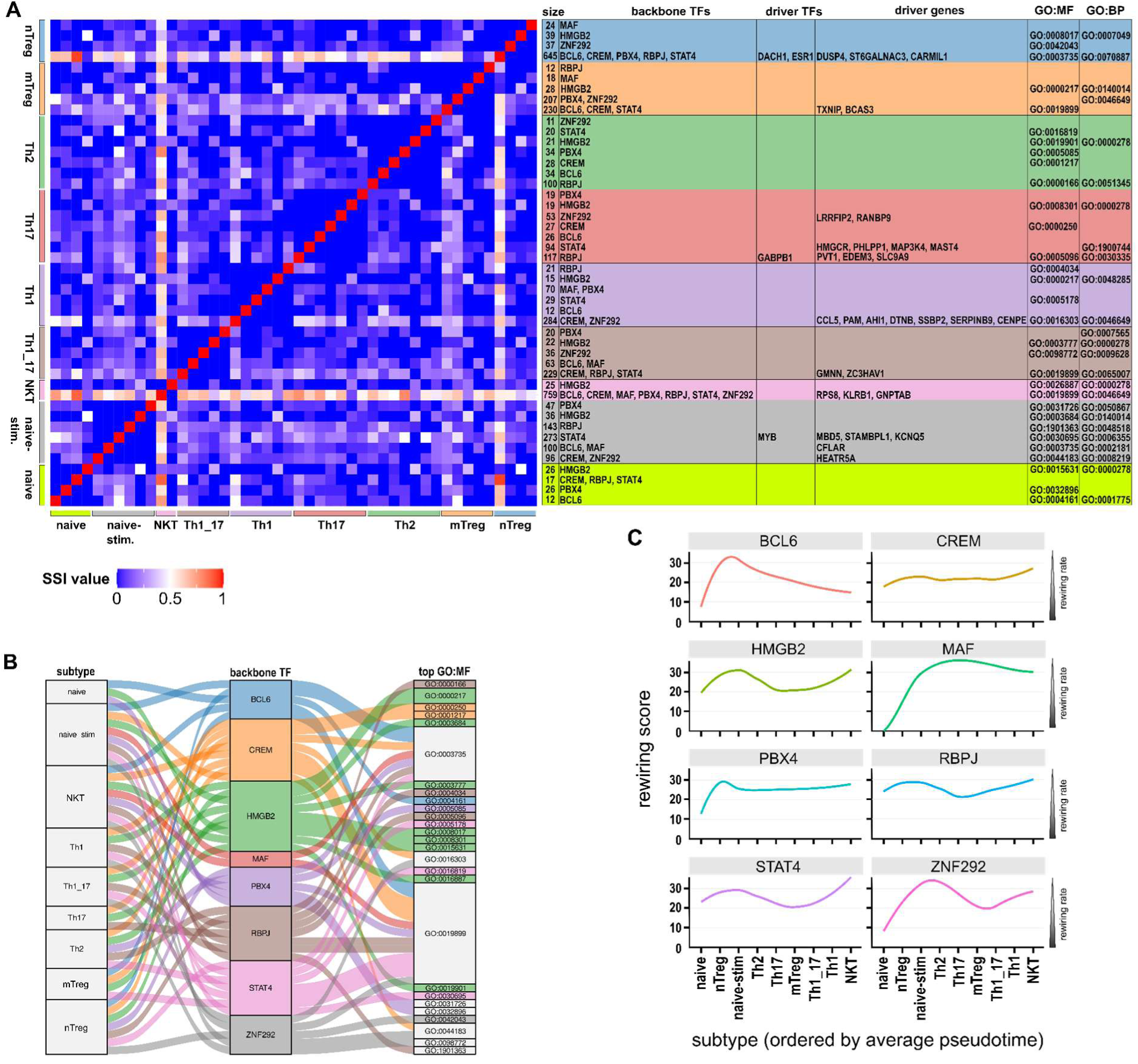
Backbone-centric community detection and backbone-specific rewiring score. (A) SSI coefficient for backbone and driver TFs and driver gene-containing communities, with the top GO:MF and GO:BP. (B) Alluvial plot with subtype, backbone-centric communities and their top GO:MF (pval < 0.05) (C) The rewiring score per pseudotime ordered-subtypes.

**Table 2:** Driver TF-containing communities. (nd = not detected)

| driver TF | subtype community | backbone TF | community size | community composition |
| --- | --- | --- | --- | --- |
| EOMES | Th1 | nd | 13 | RERE, CARD14, GTDC1, AFDN, CDK14, CDK6, FBXL20, HCN2, LIN52, MTRNR2L8, PSTPIP2, UBB |
|  | Th1_17 | nd | 12 | TPD52, PLEKHA5, ABL2, AMN1, HIST1H2BD, NIM1K, NMB, PCNA, PRKCE, TPX2, UBE2S |
| KLF2 | nd | nd | nd | nd |
| HLF | nd | nd | nd | nd |
| ZNF732 | NKT | nd |  | TANGO6, RPL19, PATJ, USP45, MAP2K6, CTNNA3, FCGR3A, MTHFD1L, SORBS1, SUGCT, NIPAL2, LGR4, CARD14, SCD5 |
| SOX13 | mTreg | nd | 15 | PPARG, AHCYL2, GCNT2, XCL2, NCAPH, CD86, PALLD, MGAT5, DMGDH, CADM1, CHN1, MAST4, TMEM65, TSC22D2 |
| TFEC | nd | nd | nd | nd |
| ESR1,<br>DACH1 | nTreg | STAT4, RBPJ,<br>PBX4, BCL6,<br>CREM | 645 | see Table S4 |
| MYB | naive | BCL6 | 12 | BCL6, CCL5, FBXL17, FDPS, ITGA4, PLIN2, SIRPG, SP140,<br>TMEM156, ARHGAP10, ACTG1 |
|  | naive-stim. | STAT4 | 273 | see Table S4 |

Overall, the SSI values across all backbone TF-centric communities were rather low, indicating substantial differences in their composition and thus confirming the rewiring (Fig. 5A). Surprisingly, the communities containing *HMGB2* seem to be the most dynamic, in terms of rewiring partners, since the SSI values tend to be the lowest observed. This is in direct opposition with FDN analysis, where *HMGB2* proved to be the most conserved backbone TF (Fig. 4C). Nevertheless, in both HMGB2 FDNs and HMGB2-containing communities, GO:BP analyses revealed similar functional involvement in the cell cycle machinery (Fig. 5A). Investigating the opposite, high SSI values spanning across most of the comparisons were documented for two communities: (1) nTreg-derived community (including *STAT4, RBPJ, PBX4, BCL6 and CREM*), and (2) NKT-derived community (including CREM, BCL6, MAF, PBX4, RBPJ, STAT4 and ZNF292). We address this observation as a potential bias of SSI metric, where both of the above-mentioned communities are proportionally larger than others (645 and 759 members, respectively), therefore the high SSI values represent only a small fraction of all the community members. Indeed, the top-ranked GO:MF/GO:BP terms annotated for individual communities seem to be highly variable with the only exception of NKT-, Th1-, Th1_17-, and mTreg-derived communities sharing either BP of “leukocyte activation” or MF of “enzyme binding”. Interestingly, CREM and STAT4 were found as conserved for “enzyme binding” and ZNF292 for “leukocyte activation” (Fig. 5A). Direct effect of different community composition is further visible on the list of high-ranked GO:MF terms - documented differences among individual communities hints at a deep rewiring whilst still maintaining high centrality (Fig. 5B).

Finally, in order to model rewiring as a function of T cell activation and differentiation, we integrated the findings of the backbone TF FDNs and community composition with pseudotime. The rewiring score for backbone TF is calculated as the product of degree centrality and the community SSI values. In this context, ‘rewiring’ refers to the alterations in gene interactions and functional associations over time. The relationship between the rewiring score and pseudotime is graphically represented, with pseudotime serving as the x-axis. This approach allows for a temporal analysis of gene network dynamics, highlighting how a gene’s connectivity and role within the network evolve over the course of the observed pseudotime (Fig.5C). We identify MAF, CREM, and RBPJ as having a low global rewiring tendency with a relatively constant degree of centrality. More dynamic changes are visible for BCL6, PBX4, HMGB2, ZNF292 in response to activation and differentiation. HMGB2 and ZNF292 show similar post-activation linked dynamics, hinting at possible synergy between these two TFs.

## Discussion

In this study, we performed an integrated single-cell analysis of genome-wide chromatin accessibility and the transcriptome to characterize the underlying differences between *in vitro* activated CD4+ T cell subtypes (Th1, Th2, Th17, Treg) and CD4+ naive cells. Additionally CD3+/CD4+ NKT cells were identified within the data. Furthermore, following a similar approach for community detection as in other studies^8,9,56–58^ and thus leveraging the resolution of multimodal scRNA-seq and scATAC-seq, we have identified central genes in the activation and differentiation of human primary CD4+ T cells in bioreactor-like conditions, as well as their rewiring patterns and community composition. To our knowledge, this is the first report to construct a multimodal-based GRN and make use of random walktrap community detection to dissect the activation and differentiation of human primary CD4+ T cells, and identify backbone genes and their associated communities. The primary analysis of multiome data was conducted to identify DE genes and DA motifs associated with DICE-annotated CD4+ T cell subsets, thus verifying the SingleR annotation and possibly identifying novel markers. Notably, a robust correlation between identified markers of DICE-annotated CD4+ T cell subsets and published data was observed, especially for distinguishing between naive and activated CD4+ T cells (Fig. 2).

Based on the subtype-wise GRN inference coupled with community detection we described the group of backbone TFs (*BCL6, CREM, HMGB2, MAF, PBX4, RBPJ, STAT4* and *ZNF292*), which showed to be highly central for the entire activation and differentiation process within all annotated CD4+ T cell subtypes. However, detailed analyses showed that despite their central role, backbone TFs’ regulatory pathways are strongly subtype-specific, which is a result of the rewiring processes. The rewiring can be understood as a joint effect between the interaction of the backbone TFs with their immediate neighborhood (FDNs) and the rest of neighbors captured by community detection (second, third, fourth, etc.). More in detail, this interaction can be of regulatory (regulator TF) or inductive (target) nature. In general, considerably low overlaps of regulator/target FDNs were documented, especially for PBX4, confirming subtype-specific rewiring events. On the other hand, the most conserved regulator TFs were found to be the HMGB1 and HIVEP2 in the case of HMGB2, the most conserved backbone TF in terms of its ubiquitous BP function as a cell cycle regulator among all subtypes (Fig. 5A). Truly, HMGB2 was shown to be important for proper differentiation and proliferation of murine CD8+ T cells^37^, which supports its similar role in CD4+ T cells as well. Similarly to the HMGB2 regulators, the first-degree targets of HMGB2 seem to be more conserved than for other backbone TFs as well (Fig. 4C). On the contrary, the composition of HMGB2-containing communities showed to be the least conserved (Fig. 5A). Nevertheless, the cellular function remains the same, hinting that despite having different community and first degree neighbor dynamics as a result of rewiring, the biological process might potentially result in a very similar outcome. Furthermore, our GRN models showed that some backbone TFs act as regulator TFs for other backbone TFs and vice versa. Investigating this, an interesting co-regulatory loop was uncovered between CREM and ZNF292. Our rewiring score captured the rewiring dynamics of the backbone TFs, showing the activation as a highly entropic/dynamic process.

MAF seems to be relevant for pro-inflammatory related subtypes, primarily Th1, Th1_17 and NKT, acting primarily in a post-activation fashion, similar to CREM and ZNF292, with a low rewiring/high conserved score. c-MAF protein has been reported to act as central repressor for IFN-y in Th1 acting downstream of IL-4 signaling, and also target IL2. Conversely, when c-MAF was depleted, increased IFN-y and Tbx21 could be observed in murine malaria and autoimmune models, in Th1 phenotype, and has a dominant role in controlling the balance between Th17 and Treg phenotypes^59,60^. c-Maf has also been shown to be critical in correct functioning of il17-producing iNKT^61^. This potentially implies the role of MAF as a rheostat-like control gene, critical for anti-inflammatory phenotype development of human primary CD4+ T cells.

Except for backbone TFs being highly central in all subtypes, 11 subtype-specific driver TFs were identified (Table 1). Among these, *EOMES* and *HLF* were suggested as important players in the Th17-Th1_17-Th1 axis. First, our scATAC data showed that EOMES motifs are highly active in Th1 and Th1_17 cells, whereas HLF motifs were active in Th1_17 and Th17 (Fig. 2). Second, both TFs were identified as highly central drivers in Th1/Th1_17 GRN (EOMES) and Th1:17/Th17 GRN (HLF)(Fig.3). These findings suggest a regulatory gradient of HLF-> EOMES TFs for Th17-> Th1 transition, which is a well-documented phenomenon^62,63^. Notably, EOMES reached the highest activity in the NKT subset (Fig. 2), however without being highly central in NKT GRN. Nevertheless, EOMES’ high activity in NKTs correlates with recent findings of its high importance within this subtype^33^.

*ESR1* and *DACH1* were identified as nTreg driver TFs with high specificity and centrality (Fig. 3). Both were also identified as members of the main nTreg community containing 5 out 8 backbone TFs and all nTreg driver genes (Fig. 5A), suggesting direct regulatory involvement of both *ESR1* and *DACH1*, in nTreg maintenance. Indeed, ESR1 was shown to be a potent regulator of CD4+ T cell differentiation with multiple roles, where in case of Tregs, ESR1 was suggested to inhibit expression of FOXP3 in the absence of estrogen^64–67^. DACH1 was shown to act as a TF negatively regulating expression of cell-cycle genes and as a tumor suppressor gene^68^, however, no direct links concerning CD4+ T cells have been described. A detailed search in nTreg GRN hasn’t shown any direct (first-degree) connections between *ESR1* and *DACH1*, nevertheless, we believe that both TFs co-orchestrate Treg development together, since both are present in the same community (Fig.5A). *SOX13* was identified as a driver TF with exclusive specificity to mTregs (Fig. 3). Only scarce information is available for *SOX13*’s role in CD4+ T cell development - *SOX13* was shown to be the key player in an early T cell development regulating αβ and γδ T cell differentiation via modifying TCF7 activity^69^. Our data also suggest a possible link for SOX13 regulating TCF7 activity in mTregs. TCF7, designated as a naive marker in our dataset, reached high expression as well as high motif activity in naive CD4+ T cells, but remarkably low expression and motif activity in mTregs (Fig. 2). Additionally, when TCF7 target FDN composition was compared between naive cells and mTregs, none of the genes were found in both groups. Thus we hypothesize that SOX13 may negatively affect TCF7 activity in mTregs. We further tried to dissect the possible roles of SOX13 in mTregs, however, *SOX13* FDN (n=14) analysis using GO:BP annotation revealed “regulation of alpha-beta T cell differentiation” (GO:0046637; p=1.081E-2) and no GO:BP terms were enriched for the *SOX13* mTreg community (Table 2; n=15). We conclude that more experiments are needed to further describe the potential role of SOX13 in mTreg development.

Finally, whilst our analysis shows a lot of transcription factors involved in known signaling pathways (STAT4, HMGB2, BCL6 etc..), none of the canonical TF (TBX21, GATA3, RORC and FOXP3) has been shown to act as highly central. This, combined with their often limited mRNA expression and enriched motifs, further reinforces the idea that the traditional “profiling” of T cells is limited in terms of defining the true identities based on protein expression only, and more attention should be focused at the transcriptomic and genomic profiles.

## Acknowledgements

The authors would like to acknowledge Dr. Johan Henriksson for designing the study, providing financial support for the data generation, as well as providing the lab equipment and space for performing the experiments.

## Funding

The computations were enabled by resources provided by the National Academic Infrastructure for Supercomputing in Sweden (NAISS) at Uppsala Multidisciplinary Center for Advanced Computational Science (UPPMAX) partially funded by the Swedish Research Council through grant agreement no. 2022-06725. I.S.M was supported by the Umeå industrial doctoral school (Företagsforskarskolan) of Umeå Universitet, and Sartorius. M.S was supported by Kempestiftelsen SMK-1959.2

## Conflicts of interest

J.T. is employed at Sartorius. I.S.M was employed at Umea university but partially funded by Sartorius, currently employed at ETH Zürich (Toxicology Lab)

## Code & Data availability

Raw sequencing data (FASTQ files) and processed single-cell multiome matrices (scRNA-seq and scATAC-seq) generated in this study will be deposited in the NCBI Gene Expression Omnibus (GEO) under accession number GSE311045 upon publication. These datasets include 10x Genomics Multiome ATAC+GEX libraries generated from human naive CD3+ CD4+ T cells differentiated under Th1, Th2, Th17, and Treg cytokine conditions. Raw sequencing runs will also be available through the Sequence Read Archive (SRA) under BioProject PRJNA1368583.

All analysis code used for preprocessing, quality control, clustering, differential accessibility/expression, integration, and figure generation is available upon reasonable request.

## Methods

### T cell isolation and culture

Blood samples were obtained from four human healthy male adults, with a mean age of 26 (ages between 20-38). Peripheral blood mononuclear cells (PBMCs) were isolated by gradient density centrifugation, using Ficoll-Paque PLUS (Cytiva, 17144002). Naïve CD4+ T cells were isolated using EasySep™ Human Naïve CD4+ T Cell Isolation Kit (Stemcell), according to manufacturer’s indications.

Human naïve CD4+ T cells were seeded in tissue culturing 96 well plates, and stimulated with plate bound anti-human CD3 Antibody (clone OKT3, 5 μg/mL, 317326, Biolegend) and aCD28 (clone CD28.2., 3 μg/mL, 302934, Biologened) at a density of 1 x 10^6/mL. The cells were activated and differentiated in ImmunoCult-XF T Cell Expansion Medium, for 5 days. The list of cytokines and concentrations used for each subtype can be found in Table S1. Activation and differentiation stimuli were added at the same time.

### Flow cytometry

On day 5, live cells were purified using flow cytometry. Cells were stained for viability discrimination using PI (P3566, Thermo Fisher Scientific), and were sorted for downstream purposes. Viability (92% avg.) was then re-assessed by Trypan Blue staining (15250061, Thermo Fisher Scientific).

### Validation of T cell differentiation

#### Flow cytometry

Human naive CD4+ T cells from two different donors were seeded on a 96-well plate (1×10^6^ cells/ml, 0.2 ml/well) and activated in Th1/Th2/Th17/Treg-differentiating conditions as described above and harvested on day 5 post activation. Non-activated human naive CD4+ T cells were cultured for 5 days as well as an activation control. For the intracellular staining of TBX21 (Th1), GATA3 (Th2), FOXP3 (Treg), and RORC (Th17) the Foxp3/Transcription Factor Staining Buffer Set (00-5523, Thermo Fisher Scientific) was used according to the manufacturer’s instructions. Following antibodies were used for the labeling of aforementioned transcription factors: Human T-bet/TBX21 Alexa Fluor® 488-conjugated Antibody (1:20, IC53851G, R&D Systems), Gata-3 Monoclonal Antibody (TWAJ), PE-Cyanine7 (1:20, 25-9966-42, Thermo Fisher Scientific), Human/Mouse ROR gamma t/RORC2/NR1F3 PE-conjugated Antibody (1:10, IC6006P, R&D Systems), and Human/Mouse/Rat FoxP3 PE-conjugated Antibody (1:10, IC8970P, R&D Systems). A BD FACSMelody Cell Sorter was subsequently used for the detection of the respective transcription factor. In case of NKT detection, freshly isolated naive CD4+ T cells from 3 donors were analyzed for the presence of CD3, CD4, and Vα24-Jα18. Following antibodies were used for the extracellular staining according to the manufacturer’s protocol: PerCP/Cyanine5.5 anti-human CD3 Antibody (#300430, BioLegend), Brilliant Violet 421™ anti-human CD4 Antibody (#357424, BioLegend), PE anti-human TCR Vα24-Jα18 (iNKT cell) Antibody (#342904, BioLegend). A BD FACSMelody Cell Sorter was subsequently used for the detection of CD3+/CD4+/Vα24-Jα18+ cells.

### Single-cell multiomics library preparation and sequencing

Dissociated cells were washed in ice-cold ATAC-seq resuspension buffer (RSB, 10 mM Tris pH 7.4, 10 mM NaCl, 3 mM MgCl2), spun down, and resuspended in 100 mL ATAC-seq lysis buffer (RSB plus 0.1% NP-40 and 0.1% Tween-20 (Thermo Fisher). Lysis was allowed to proceed on ice for 5 min, then 900 mL RSB was added before spinning down again and resuspending in 50 mL 1X Nuclei Resuspension Buffer (10x Genomics). To assess nuclei purity and integrity after lysis, nuclei were stained with Trypan Blue (15250061, Thermo Fisher Scientific), and DAPI (D1306, Thermo Fisher Scientific), according to manufacturer recommendation. If necessary, cell concentrations were adjusted to equal ratios prior to starting single-cell GEM emulsion droplet generation with the ATAC-seq NextGEM kit (10x Genomics). Briefly, nuclei were incubated in a transposition mix. Transposed nuclei were then loaded into a Chromium Next GEM Chip J. 9,000 nuclei were loaded per lane, with a target recovery of 5,500 (doublet rate 4% - 4.8%). After GEM emulsion generation, reverse transcription, cDNA amplification and multiome sc library generation occurred, according to manufacturer specifications (https://assets.ctfassets.net/an68im79xiti/5tnYerYC4FdDwYK0hlcfWj/cc46120ef02327249b5c42f24f44d699/CG000338_ChromiumNextGEM_Multiome_ATAC_GEX_User_Guide_RevA.pdf)

### Multiome Single-cell preprocessing

Library reads were aligned and aggregated using CellRanger ARC 2.0.0. Single-cell RNA-seq analysis was done using Seurat^19^, and single-cell ATAC-seq analysis using Signac^20^ following the standard pipeline. Briefly, after initial QC filtering of scRNA-seq data (mtDNA content < 7.5%, 200 < genes/cell < 7000, 1000 < reads/cell < 25 000) the cell cycle scoring was performed using the list of CC-related genes^70^. SCTransform() was used for normalization, finding variable features and scaling. The cell cycle regression was not performed, since PCA did not reveal any separation of cells based on their cell cycle scoring. The clustering and dimensionality reduction was computed using PCA-based UMAP^21^ (dims=1:30, resolution = 0.5). Cell types were annotated using SingleR automated annotation^22^ by employing the DICE reference dataset^23^. Cell types, making up less than 150 cells/category were removed from the analysis (naive B cells [7 cells], monocytes CD16+ [1 cell], monocytes CD14+ [2 cells], CD8+ naive T cells [44 cells], CD4+ TFH cells [89], CD8+ naive-stimulated T cells [124 cells]). In case of scATAC-seq data the QC filtering (1000 < reads/cell < 100 000, nucleosome signal < 2, TSS enrichment score >1) was followed by peak calling using MACS2^71^, which was grouped by “SingleR labels” (i.e. annotated cell types). The clustering and dimensionality reduction was computed using LSI-based UMAP^21^ (nNeighbors = 30, dims=2:30, resolution = 0.7). Subsequently, joint UMAP visualization for both, scRNA-seq and scATAC-seq, was performed using PCA and LSI reductions (dims=1:30 and 2:30, respectively). Phylogenetic analysis of DICE-annotated T cell subsets was performed by employing BuildClusterTree(); PCA reduction (dims=1:30) and LSI reduction (dims=2:30) was used for the analysis in case of scRNA-seq and scATAC-seq data, respectively. For the identification of marker genes, each T cell subset was compared separately to CD4+ T naive cells via FindMarkers() function. Subsequent GSEA (Gene set enrichment analysis) was performed using clusterProfiler^72^ and function gseGO(ont=ALL, pAdjustMethod=BH, pvalueCutoff=0.05).

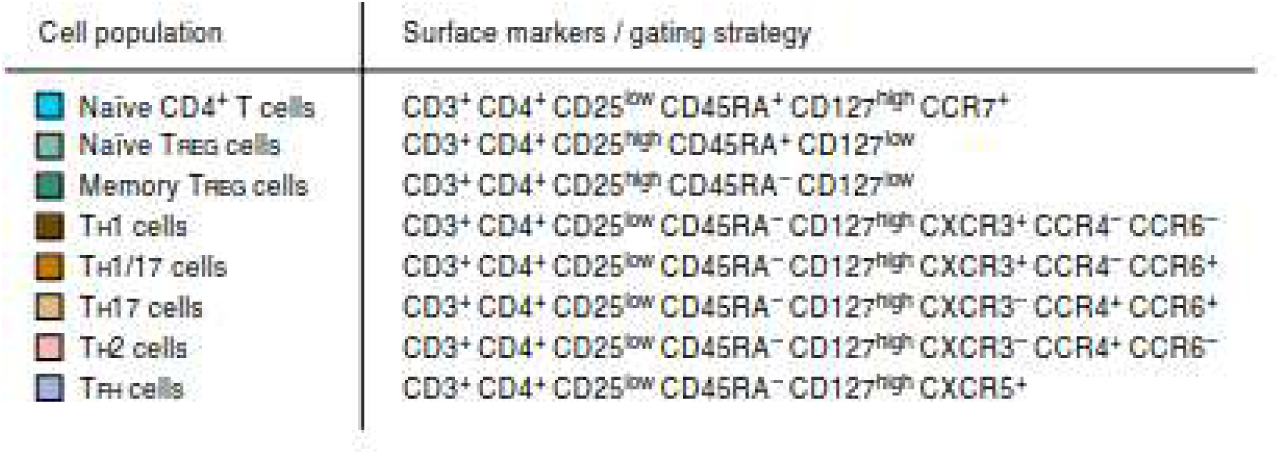

### Network construction using Pando

Pando creates a putative GRN by finding TF-DNA binding site pairs, and then infers a TF-target GRN using one of several regression models implementation, for each single cell^4^. XGBOOST was used for network inference.Then, Find_modules (pval<0.05, rsq threshold= 0.1, max 50 targets per module) and Get_network_graph embedding. Networks were validated by comparing the means of the degree centrality distributions of the predicted networks with 1000 random networks and setting a pval< 0.05. The simulated graphs were generated using Igraph’s watts.strogatz.game function, taking into account nr of nodes, average number of FDNs and the average edge density of the predicted networks.

### Network and community analysis

Networks were aggregated using igraph’s graph.intersection function, which keeps identical or overlapping vertex sets. Backbone and Driver TFs and Driver genes were identified using degree and eigen.centrality functions from igraph. FDNs were retrieved using igraph’s degree. Pairwise overlap was determined by using the node Szymkiewich-Simpson coefficient (SSI) *SSI* = *N*(*A*) ∩ *N*(*B*) / *min*(∣ *N*(*A*) ∣, ∣ *N*(*B*) ∣) ∣ where A and B are two different graphs Communities were identified using cluster_walktrap with 4 steps and no weights, from the igraph package. GSEA analysis was performed in Gprofiler (cite) using GO:BP or GO:MF databases, with annotated functions, FDR correction method, and a significance score of < 0.05, for homo sapiens.

### Rewiring score

The rewiring score per backbone was calculated as *RS* = *DC*(*x*, *y*) ∗ *SSI*(*x*, *z*, *y*), where DC is degree centrality of backbone X in network Y, and SSI is Szymkiewicz-Simpson coefficient of backbone X, in community Z and in network Y.

## Notes

### Competing Interest Statement

J.T. is employed at Sartorius. I.S.M was employed at Umea university but partially funded by Sartorius. Other authors declare no conflict of interest.

## References

1. Soskic, B. et al. Immune disease risk variants regulate gene expression dynamics during CD4+ T cell activation. Nat. Genet. 54, 817–826 (2022).

2. Cano-Gamez, E. et al. Single-cell transcriptomics identifies an effectorness gradient shaping the response of CD4+ T cells to cytokines. Nat. Commun. 11, 1–15 (2020).

3. Karlebach, G. & Shamir, R. Modelling and analysis of gene regulatory networks. Nat. Rev. Mol. Cell Biol. 9, 770–780 (2008).

4. Fleck, J. S. et al. Inferring and perturbing cell fate regulomes in human brain organoids. Nature 1–8 (2022).

5. Kamimoto, K. et al. Dissecting cell identity via network inference and in silico gene perturbation. Nature 614, 742–751 (2023).

6. Calderer, G. & Kuijjer, M. L. Community Detection in Large-Scale Bipartite Biological Networks. Front. Genet. 12, (2021).

7. Website. https://www.pnas.org/doi/abs/10.1073/pnas.0601602103.

8. Zhang, K. et al. Longitudinal single-cell RNA-seq analysis reveals stress-promoted chemoresistance in metastatic ovarian cancer. Sci Adv 8, eabm1831 (2022).

9. Song, W.-M. et al. Network models of primary melanoma microenvironments identify key melanoma regulators underlying prognosis. Nat. Commun. 12, 1–14 (2021).

10. Eldar, A. & Elowitz, M. B. Functional roles for noise in genetic circuits. Nature 467, 167–173 (2010).

11. Raj, A. & van Oudenaarden, A. Nature, nurture, or chance: stochastic gene expression and its consequences. Cell 135, (2008).

12. Delvenne, J.-C., Yaliraki, S. N. & Barahona, M. Stability of graph communities across time scales. Proceedings of the National Academy of Sciences 107, 12755–12760 (2010).

13. Rosvall, M. & Bergstrom, C. T. Maps of random walks on complex networks reveal community structure. Proceedings of the National Academy of Sciences 105, 1118–1123 (2008).

14. Community detection in graphs. Physics Reports 486, 75–174 (2010).

15. Oguchi, A. et al. An atlas of transcribed enhancers across helper T cell diversity for decoding human diseases. Science (2024) doi:10.1126/science.add8394.

16. King, H. W. et al. Integrated single-cell transcriptomics and epigenomics reveals strong germinal center–associated etiology of autoimmune risk loci. Science Immunology (2021) doi:10.1126/sciimmunol.abh3768.

17. Mitra, S. et al. Single-cell multi-ome regression models identify functional and disease-associated enhancers and enable chromatin potential analysis. Nature Genetics 56, 627–636 (2024).

18. Chopp, L. B. et al. An Integrated Epigenomic and Transcriptomic Map of Mouse and Human αβ T Cell Development. Immunity 53, 1182–1201.e8 (2020).

19. Hao, Y. et al. Integrated analysis of multimodal single-cell data. Cell 184, 3573–3587.e29 (2021).

20. Stuart, T., Srivastava, A., Madad, S., Lareau, C. A. & Satija, R. Single-cell chromatin state analysis with Signac. Nat. Methods 18, 1333–1341 (2021).

21. McInnes, L., Healy, J., Saul, N. & Großberger, L. UMAP: Uniform Manifold Approximation and Projection. J. Open Source Softw. 3, 861 (2018).

22. Aran, D. et al. Reference-based analysis of lung single-cell sequencing reveals a transitional profibrotic macrophage. Nat. Immunol. 20, 163–172 (2019).

23. Schmiedel, B. J. et al. Impact of Genetic Polymorphisms on Human Immune Cell Gene Expression. Cell 175, 1701–1715.e16 (2018).

24. Krijgsman, D., Hokland, M. & Kuppen, P. J. K. The Role of Natural Killer T Cells in Cancer—A Phenotypical and Functional Approach. Front. Immunol. 9, (2018).

25. Nakagawa, R., Brayer, J., Restrepo, N., Mulé, J. J. & Mailloux, A. W. High-Dimensional Flow Cytometry Analysis of Regulatory Receptors on Human T Cells, NK Cells, and NKT Cells. Methods Mol. Biol. 2194, 255–290 (2021).

26. Schep, A. N., Wu, B., Buenrostro, J. D. & Greenleaf, W. J. chromVAR: inferring transcription-factor-associated accessibility from single-cell epigenomic data. Nat. Methods 14, 975–978 (2017).

27. Atsaves, V., Leventaki, V., Rassidakis, G. Z. & Claret, F. X. AP-1 Transcription Factors as Regulators of Immune Responses in Cancer. Cancers 11, (2019).

28. McKarns, S. C. & Schwartz, R. H. Distinct effects of TGF-β1 on CD4+ and CD8+ T cell survival, division, and IL-2 production: a role for T cell intrinsic Smad3. The Journal of Immunology (2005).

29. Ivashkiv, L. B. IFNγ: signalling, epigenetics and roles in immunity, metabolism, disease and cancer immunotherapy. Nat. Rev. Immunol. 18, 545–558 (2018).

30. Szabo, S. J. et al. A novel transcription factor, T-bet, directs Th1 lineage commitment. Cell 100, 655–669 (2000).

31. Dejean, A. S., Joulia, E. & Walzer, T. The role of Eomes in human CD4 T cell differentiation: A question of context. Eur. J. Immunol. 49, 38–41 (2019).

32. Mazzoni, A. et al. Eomes controls the development of Th17-derived (non-classic) Th1 cells during chronic inflammation. Eur. J. Immunol. 49, 79–95 (2019).

33. Shimizu, K. et al. Eomes transcription factor is required for the development and differentiation of invariant NKT cells. Commun Biol 2, 150 (2019).

34. Shui, X. et al. Knockdown of lncRNA NEAT1 inhibits Th17/CD4+ T cell differentiation through reducing the STAT3 protein level. J. Cell. Physiol. 234, 22477–22484 (2019).

35. Ivanov, I. I. et al. The Orphan Nuclear Receptor RORγt Directs the Differentiation Program of Proinflammatory IL-17+ T Helper Cells. Cell 126, 1121–1133 (2006).

36. Luo, Y. et al. Single-cell transcriptomic analysis reveals disparate effector differentiation pathways in human Treg compartment. Nat. Commun. 12, 3913 (2021).

37. Neubert, E. N. et al. HMGB2 regulates the differentiation and stemness of exhausted CD8+ T cells during chronic viral infection and cancer. Nat. Commun. 14, 5631 (2023).

38. Tsukumo, S.-I. et al. Bach2 maintains T cells in a naive state by suppressing effector memory-related genes. Proc. Natl. Acad. Sci. U. S. A. 110, 10735–10740 (2013).

39. Odagiu, L., May, J., Boulet, S., Baldwin, T. A. & Labrecque, N. Role of the Orphan Nuclear Receptor NR4A Family in T-Cell Biology. Front. Endocrinol. 11, (2021).

40. Zhang, Y. et al. Impaired apoptosis, extended duration of immune responses, and a lupus-like autoimmune disease in IEX-1-transgenic mice. Proc. Natl. Acad. Sci. U. S. A. 99, 878–883 (2002).

41. Kim, H.-J. et al. Stable inhibitory activity of regulatory T cells requires the transcription factor Helios. Science 350, 334–339 (2015).

42. Miyara, M. et al. Functional delineation and differentiation dynamics of human CD4+ T cells expressing the FoxP3 transcription factor. Immunity 30, 899–911 (2009).

43. Escobar, G., Mangani, D. & Anderson, A. C. T cell factor 1: A master regulator of the T cell response in disease. Sci Immunol 5, (2020).

44. Willinger, T. et al. Human naive CD8 T cells down-regulate expression of the WNT pathway transcription factors lymphoid enhancer binding factor 1 and transcription factor 7 (T cell factor-1) following antigen encounter in vitro and in vivo. J. Immunol. 176, 1439–1446 (2006).

45. Garaud, S. et al. FOXP1 is a regulator of quiescence in healthy human CD4+ T cells and is constitutively repressed in T cells from patients with lymphoproliferative disorders. Eur. J. Immunol. 47, 168–179 (2017).

46. Dynamic regulatory networks of T cell trajectory dissect transcriptional control of T cell state transition. Molecular Therapy - Nucleic Acids 26, 1115–1129 (2021).

47. Barabasi, A. L. & Albert, R. Emergence of scaling in random networks. Science 286, 509–512 (1999).

48. Choi, J. & Crotty, S. Bcl6-Mediated Transcriptional Regulation of Follicular Helper T cells (TFH). Trends Immunol. 42, 336–349 (2021).

49. Verjans, E. et al. The cAMP response element modulator (CREM) regulates TH2 mediated inflammation. Oncotarget 6, 38538–38551 (2015).

50. Lippe, R. et al. CREMα overexpression decreases IL-2 production, induces a T(H)17 phenotype and accelerates autoimmunity. J. Mol. Cell Biol. 4, 121–123 (2012).

51. Haining, W. N. & Weiss, S. A. c-Maf in CD4+ T cells: it’s all about context. Nat. Immunol. 19, 429–431 (2018).

52. Chen, E. L. Y., Thompson, P. K. & Zúñiga-Pflücker, J. C. RBPJ-dependent Notch signaling initiates the T cell program in a subset of thymus-seeding progenitors. Nat. Immunol. 20, 1456–1468 (2019).

53. Delacher, M. et al. Rbpj expression in regulatory T cells is critical for restraining TH2 responses. Nat. Commun. 10, 1621 (2019).

54. O’Shea, J. J., Lahesmaa, R., Vahedi, G., Laurence, A. & Kanno, Y. Genomic views of STAT function in CD4+ T helper cell differentiation. Nat. Rev. Immunol. 11, 239–250 (2011).

55. Website. 10.48550/arXiv.physics/0512106 doi:10.48550/arXiv.physics/0512106.

56. Vrahatis, A. G., Dimitrakopoulos, G. N., Tasoulis, S. K., Georgakopoulos, S. V. & Plagianakos, V. P. Single-cell regulatory network inference and clustering from high-dimensional sequencing data. in 2019 IEEE International Conference on Big Data (Big Data) (IEEE, 2019). doi:10.1109/bigdata47090.2019.9006016.

57. Website. 10.7554/eLife.56221 doi:10.7554/eLife.56221.

58. Single-cell proteomics defines the cellular heterogeneity of localized prostate cancer. Cell Reports Medicine 3, 100604 (2022).

59. Gabryšová, L. et al. c-Maf controls immune responses by regulating disease-specific gene networks and repressing IL-2 in CD4+ T cells. Nat. Immunol. 19, 497–507 (2018).

60. Imbratta, C., Hussein, H., Andris, F. & Verdeil, G. c-MAF, a Swiss Army Knife for Tolerance in Lymphocytes. Front. Immunol. 11, (2020).

61. Yu, J.-S. et al. Differentiation of IL-17-Producing Invariant Natural Killer T Cells Requires Expression of the Transcription Factor c-Maf. Front. Immunol. 8, (2017).

62. Guéry, L. & Hugues, S. Th17 Cell Plasticity and Functions in Cancer Immunity. Biomed Res. Int. 2015, 314620 (2015).

63. Stadhouders, R., Lubberts, E. & Hendriks, R. W. A cellular and molecular view of T helper 17 cell plasticity in autoimmunity. J. Autoimmun. 87, 1–15 (2018).

64. Lélu, K. et al. Estrogen receptor α signaling in T lymphocytes is required for estradiol-mediated inhibition of Th1 and Th17 cell differentiation and protection against experimental autoimmune encephalomyelitis. J. Immunol. 187, 2386–2393 (2011).

65. Kim, D.-H. et al. Estrogen receptor α in T cells suppresses follicular helper T cell responses and prevents autoimmunity. Exp. Mol. Med. 51, 1–9 (2019).

66. Mohammad, I. et al. Estrogen receptor α contributes to T cell-mediated autoimmune inflammation by promoting T cell activation and proliferation. Sci. Signal. 11, (2018).

67. Goodman, W. A. et al. Impaired estrogen signaling underlies regulatory T cell loss-of-function in the chronically inflamed intestine. Proc. Natl. Acad. Sci. U. S. A. 117, 17166–17176 (2020).

68. Merscher, S. & Faul, C. DACH1 as a multifaceted and potentially druggable susceptibility factor for kidney disease. J. Clin. Invest. 131, (2021).

69. Melichar, H. J. et al. Regulation of gammadelta versus alphabeta T lymphocyte differentiation by the transcription factor SOX13. Science 315, 230–233 (2007).

70. Tirosh, I. et al. Dissecting the multicellular ecosystem of metastatic melanoma by single-cell RNA-seq. Science 352, 189–196 (2016).

71. Zhang, Y. et al. Model-based analysis of ChIP-Seq (MACS). Genome Biol. 9, R137 (2008).

72. Wu, T. et al. clusterProfiler 4.0: A universal enrichment tool for interpreting omics data. Innovation (Camb) 2, 100141 (2021).

