## Supplemental figures for "A conserved transcriptional backbone and rewiring of gene-regulatory networks in activated human CD4⁺ T cells"

**Affiliations:**
1. Molecular Infection Medicine Sweden (MIMS), Umeå Centre for Microbial Research, Department of Molecular Biology, Umeå University, Umeå, Sweden

2. Industrial Doctoral School, Umeå University, Umeå, Sweden.

3. Department of Clinical Microbiology, Umeå University, Umeå, Sweden

4. Barcelona Supercomputing Center, Plaça Eusebi Güell, 1-3, 08034 Barcelona, Spain

5. Computational Life Science Cluster (CLiC), Umeå University, Umeå, Sweden.

6. Sartorius Corporate Research, Umeå, Sweden.

7. Department of Health Sciences and Technology(D-HEST), ETH Zürich, Zürich, Switzerland

8. Faculty of Science, University of South Bohemia, České Budějovice, Czech Republic

* Corresponding authors
‡ These authors contributed equally to this work

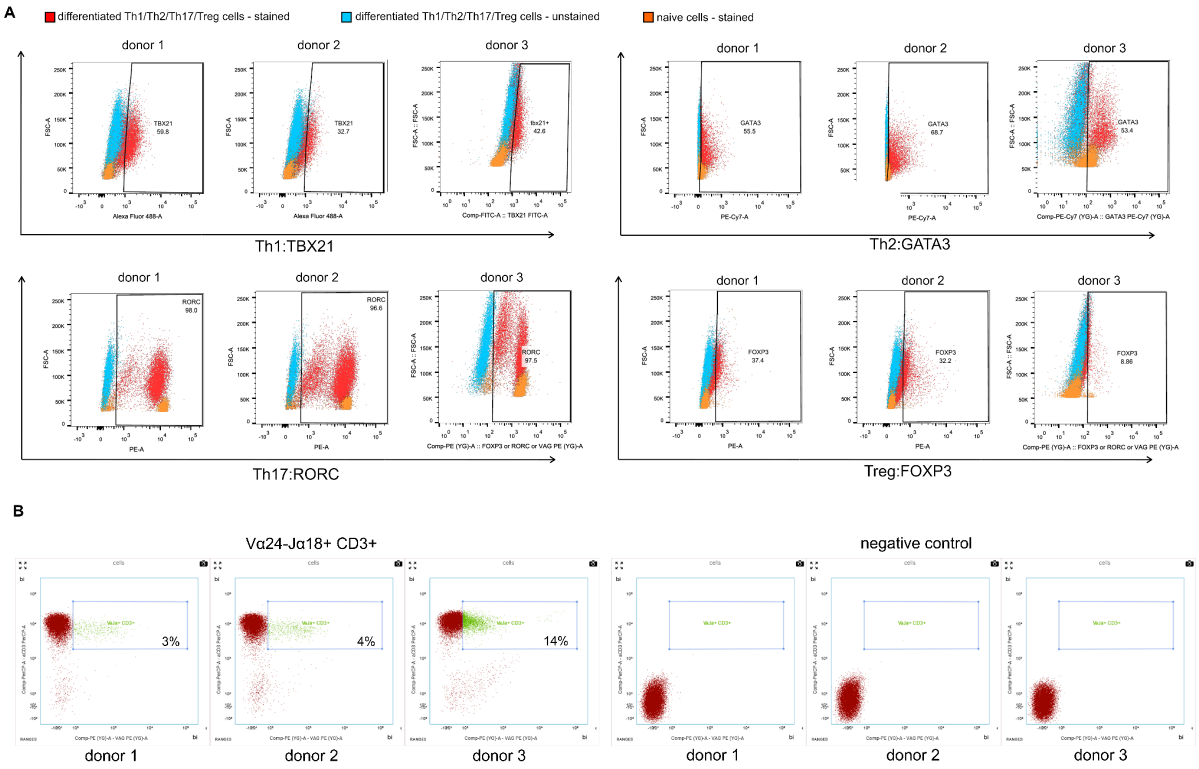


*Figure S1: (A) Flow cytometry analysis of in vitro differentiated and naive CD4+ T cells. Human naive CD4+ T cells were activated and differentiated using Th1/Th2/Th17/Treg-specific cytokine/antibody cocktails for 5 days. Differentiated and naive T cell populations were analyzed for the expression of Th1/Th2/Th17/Treg canonical markers (TBX21, GATA3, RORC, FOXP3, respectively). Each graph represents a merged plotting of stained/unstained T cells of the respective subtype. Values represent the percentage of marker-positive differentiated cells after gating on non-labeled control. (B) FACS analysis of NKT content (Vα24-Jα18+/CD3+) in freshly isolated naive CD4+ T cells from blood-derived PBMCs (n= 3 donors). note: this panel is not in its final form (these are just temporary images).*

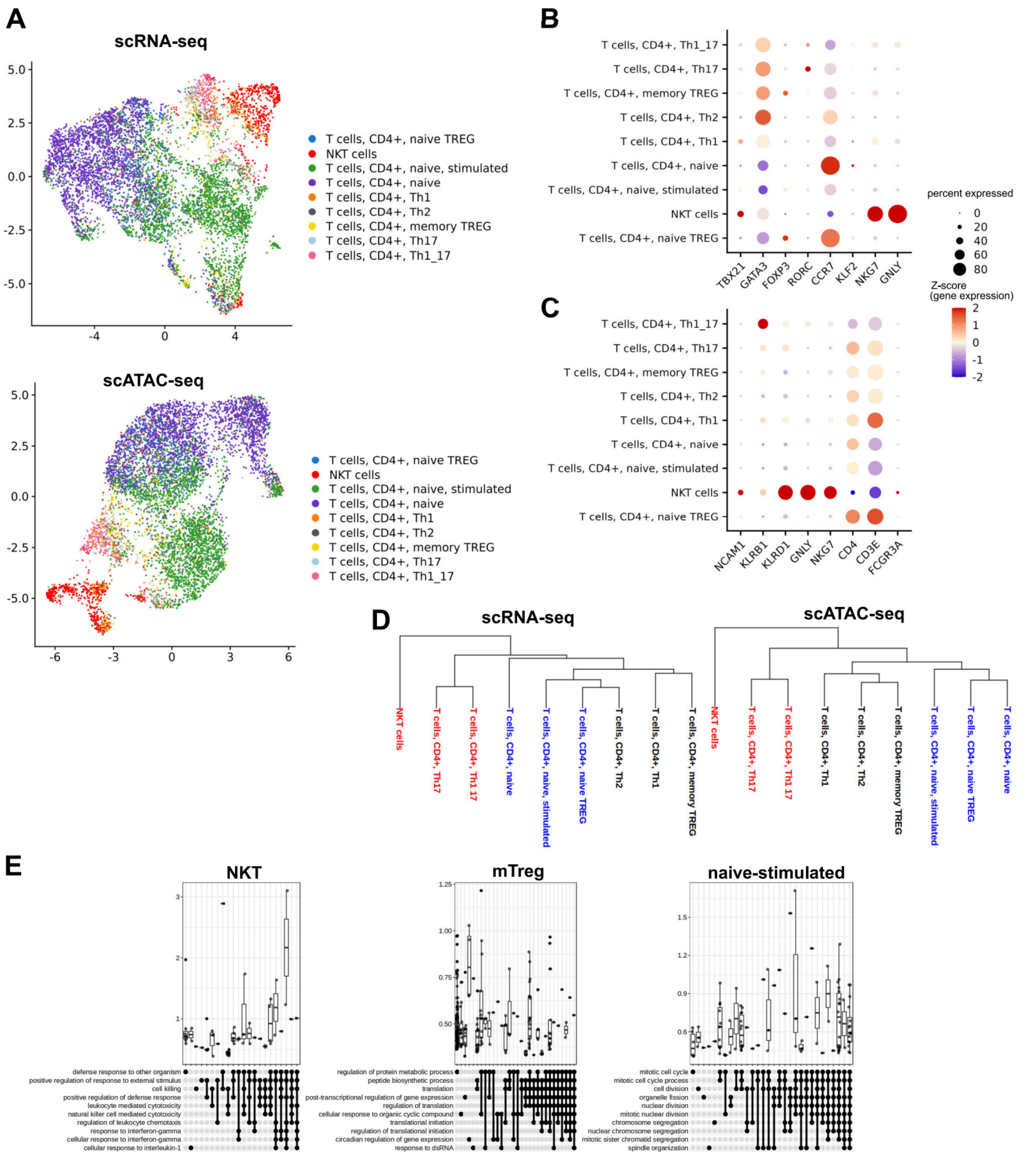


*Figure S2: Single cell RNA-seq and ATAC-seq (multiome-seq) analysis of in vitro differentiated human CD4+ T cells and manual re-annotation for NKT cells. (A) UMAP embedding for scRNA-seq data (left) and scATAC-seq data (right) with highlighted DICE-annotated T cell subtypes. (B) Dot plot representing Z-score values of average expression for selected CD4+ T cell marker genes: TBX21 (Th1), GATA3 (Th2), FOXP3 (Tregs), RORC (Th17), CCR7 and KLF2 (naive T cells), NKG7 and GNLY (NK cells). (C) Dot plot representing Z-score values of average expression for selected NK/NKT markers. The crucial feature of NKT cells is the co-expression of NK markers (KLRB1, KLRD1, GNLY, NKG7) with NCAM1 and CD3. (D) Phylogenetic clustering analysis based on scRNA-seq data (left) and scATAC-seq data (right). Color code represents two main subclusters from scATAC-seq data (black, blue) and Th1_17/Th17 subcluster combined with outgrouping NKT cells (red). (E) GSEA analysis of DE gene sets resulted in significant hit only for NKT cells, mTregs, and naive-stimulated cells (Benjamini-Hochberg adjusted $p < 0.05$).*

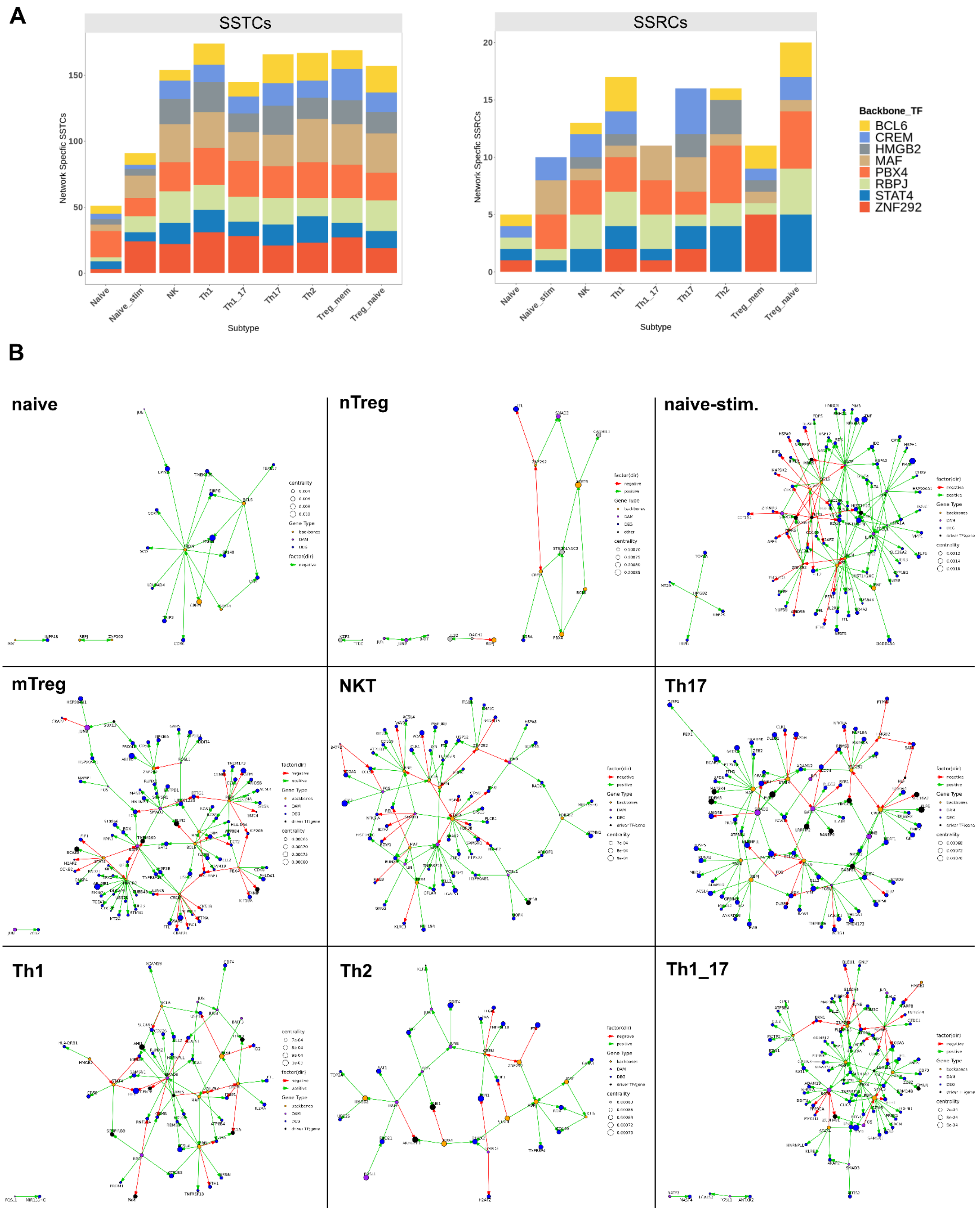


*Figure S3: GRN analysis of backbone-centric first degree neighbor edges. (A) Barplot with absolute numbers of subtype-specific target connections (SSTCs; left) and subtype-specific regulator connections (SSRCs; right). (B) Visualization of Pando-derived GRNs including only FDNs connected to backbone TFs (orange), respective DEGs (blue) and DAMs (purple) or driver TFs/genes (black).*

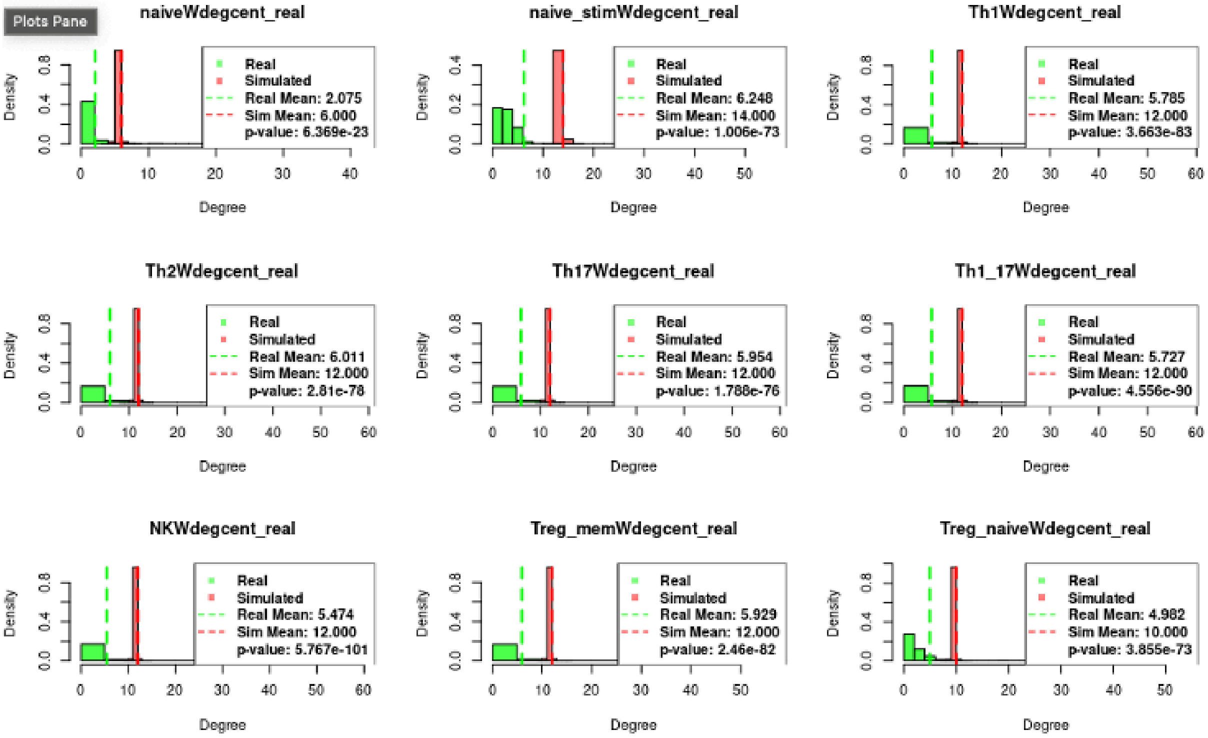


*Figure S4: Network validation. 1000 random networks were simulated using Watts Strogatz algorithm. For each simulated batch of networks similar values of the nr of nodes, average number of FDNS and average edge density were used as criteria. Means of degree centrality distributions were compared using a two tailed T test with a p.val < 0.05*
